# Distinct activation programs in naive and memory CD8 T cells govern progeny stemness and effector persistence

**DOI:** 10.64898/2026.07.22.740179

**Authors:** Eva Salyova, Darina Paprckova, Juraj Michalik, Veronika Niederlova, Veronika Cimermanova, Katerina Tomicova, Ales Drobek, Vojtech Racek, Alena Moudra, Anna Morales Mendez, Michaela Krupkova, Radislav Sedlacek, Ondrej Stepanek

## Abstract

Infection- and vaccination-induced memory CD8^+^ T cells provide protection upon subsequent exposure to cognate antigen through their increased abundance and robust per-cell responses. However, how prior antigen experience alters T-cell activation programs remains poorly understood. We longitudinally profiled gene expression in naive, central memory, and effector memory CD8^+^ T cells in response to cognate antigen across multiple models of acute infection. Naive T cells engaged TOX- and TCF7-centered programs and generated early stem-like central memory precursors. Their initially slow proliferation was followed by rapid expansion and the production of large numbers of short-lived effector cells. Central memory T cells rapidly triggered effector and proliferation programs while maintaining a smaller self-renewing population. Effector memory T cells expanded poorly and generated almost exclusively effector progeny. Effector progeny derived from both memory subsets survived contraction more efficiently than naive T-cell-derived progeny and established persistent effector memory populations. These data show that prior antigen experience does not simply accelerate CD8^+^ T-cell activation but redirects cell-intrinsic programs that shape progeny fate and persistence. These distinct programs may reflect adaptation to primary versus repeated antigen exposure and suggest that infection and vaccination history can shape the balance between stem-like memory and persistent effector populations.

## INTRODUCTION

Immune memory is one of the key features of adaptive immunity. A fraction of antigen-specific T cells generated during a primary response persists after antigen clearance and responds rapidly upon reinfection ^1, 2^. During the primary response, antigen-specific naive T cells (Tn) undergo clonal expansion and form heterogeneous progeny of effector (Teff) and memory T cells (Tm). Whereas neutralizing antibodies typically recognize viral epitopes involved in host-cell entry, protective Tm cells can recognize MHC-presented peptides derived from proteins throughout the viral proteome, including internal and non-structural proteins ^3^. Thus, Tm-mediated protection can be more resilient to antigenic variation than neutralizing antibody responses in viral infections ^4, 5^. Accordingly, there is renewed interest in developing vaccines that elicit broad and durable T-cell memory ^6, 7^.

Circulating CD8^+^ Tm cells comprise subsets with distinct migratory and functional properties ^8, 9^. Central memory T cells (Tcm) recirculate through secondary lymphoid tissues and retain substantial proliferative and self-renewal capacity, whereas effector memory T cells (Tem) are biased toward peripheral trafficking and rapid effector function ^10, 11^. In addition, antigen-inexperienced memory-like T cells (Taim), also termed virtual or innate memory T cells, acquire memory-associated properties through homeostatic signaling without prior encounter with cognate foreign antigen ^12, 13, 14^.

The protective advantage of Tm cells reflects both their increased abundance and their rapid deployment of effector functions on a per-cell basis ^15, 16^. Depending on the context, Tm-mediated protection can involve IFNγ production ^17^ and/or direct cytotoxicity ^18^. Tm cells retain epigenetically poised effector genes, including IFNG, that support rapid recall responses ^19, 20^. Moreover, Tem cells are equipped for eliciting immediate cytotoxicity ^21, 22^.

The proliferative capacity and cumulative expansion of CD8^+^ Tn and Tm cells remain unresolved. Some studies have reported similar clonal expansion of Tn and Tm cells ^11, 23, 24^, whereas others observed faster or greater proliferation or expansion of Tm cells ^15, 25^ or Tn cells ^26, 27, 28^. These apparently conflicting findings may reflect differences in antigen abundance and persistence, inflammatory context, memory-subset composition, and timing. The expansion of the memory compartment depends on its composition, since Tcm cells proliferate more rapidly upon antigenic restimulation than Tem cells ^11, 23, 29^.

Tn and Tm cells also differ in their integration of antigenic and innate signals. Some studies have reported reduced proximal TCR signaling or higher antigen requirements in Tm than in Tn cells under defined conditions ^27, 30^, whereas Tm and Taim cells respond efficiently to inflammatory cytokines and can undergo antigen-independent bystander activation ^31, 32^. Altogether, the in vivo responses of Tn and Tm cells to cognate antigen are shaped by their distinct responses to antigenic and innate signals, as well as by their pre-existing epigenetic states. Although modern transcriptomic studies have mapped differentiation trajectories arising from Tn cells during primary infection ^33, 34^, the activation-induced gene-expression programs of pre-existing Tn and Tm cells have not been systematically compared in vivo. Thus, it is unclear whether prior antigen experience merely alters the kinetics and magnitude of a conserved T-cell activation program or redirects the program itself ^1, 2^.

In this study, we longitudinally profiled the transcriptional and functional responses of CD8^+^ Tn, Tcm, Tem, and Taim cells during acute infection. We found that these populations engaged distinct cell-intrinsic activation programs that shaped the fates of their progeny. Tn and Taim cells activated programs that promoted stemness and favored the generation of Tcm progeny, whereas Tcm cells retained a degree of self-renewal but preferentially engaged an effector differentiation program, and Tem cells generated predominantly Teff progeny. In addition, Teff cells derived from Tcm and Tem cells survived better and persisted as Tem cells more efficiently than Teff cells derived from Tn cells. These findings reveal that prior antigen experience does not merely accelerate recall but reshapes cell-intrinsic activation programs that govern progeny fate and persistence.

## RESULTS

### Distinct early activation programs of Tn, Tm, and Taim CD8^+^ T cells

To compare antigenic responses of CD8^+^ Tn, Taim, and Tm cells in vivo, we generated and isolated these three T-cell populations expressing the same retrogenic K^b^-OVA-specific TCR, either clone 2 or clone 12 ^13, 35^, and transferred them to CD45.1 congenic host mice, which were subsequently infected with *Listeria monocytogenes* expressing ovalbumin (Lm-OVA) (Fig. 1A, Supplementary Fig. 1A-B). On day 3 post-infection (p.i.), we analyzed the transcriptional profile of progeny of the transferred cells in bulk. We observed the largest differences between the Tn and Tm progeny, whereas Taim progeny showed intermediate signature (Fig. 1B-C, Supplementary Fig. 1C-E). The Tm progeny upregulated effector genes (*Gzmb*, *Prf1*, *Ccl5, Klrg1*), effector transcription factors (*Zeb2*, *Id2, Prdm1*), integrins (*Itga1*, *Itgax*), and *Mt1-3* genes encoding metallothioneins, whereas Tn progeny upregulated genes associated with Tcm signature (*Sell*, *Ccr7*), transcription factors associated with stemness (*Tcf7*, *Lef1*, *Id3*, *Bach2*) (Fig. 1C). Some inhibitory receptors (*Pdcd1*, *Ctla4*, *Lag3, Slamf6*) and their regulator *Tox* were upregulated in Tn progeny cells, whereas other inhibitory receptors (*Havcr2/Tim3* and *Tigit*) were upregulated in Tm progeny cells (Fig. 1B-C). We verified differential expression of selected genes using RT-qPCR on sorted subsets (Supplementary Fig. 1F). The gene set enrichment analyses (GSEA) revealed upregulation of effector-signature and downregulation of memory-signature genes in Tm progeny (Fig. 1D).

**Fig. 1.**
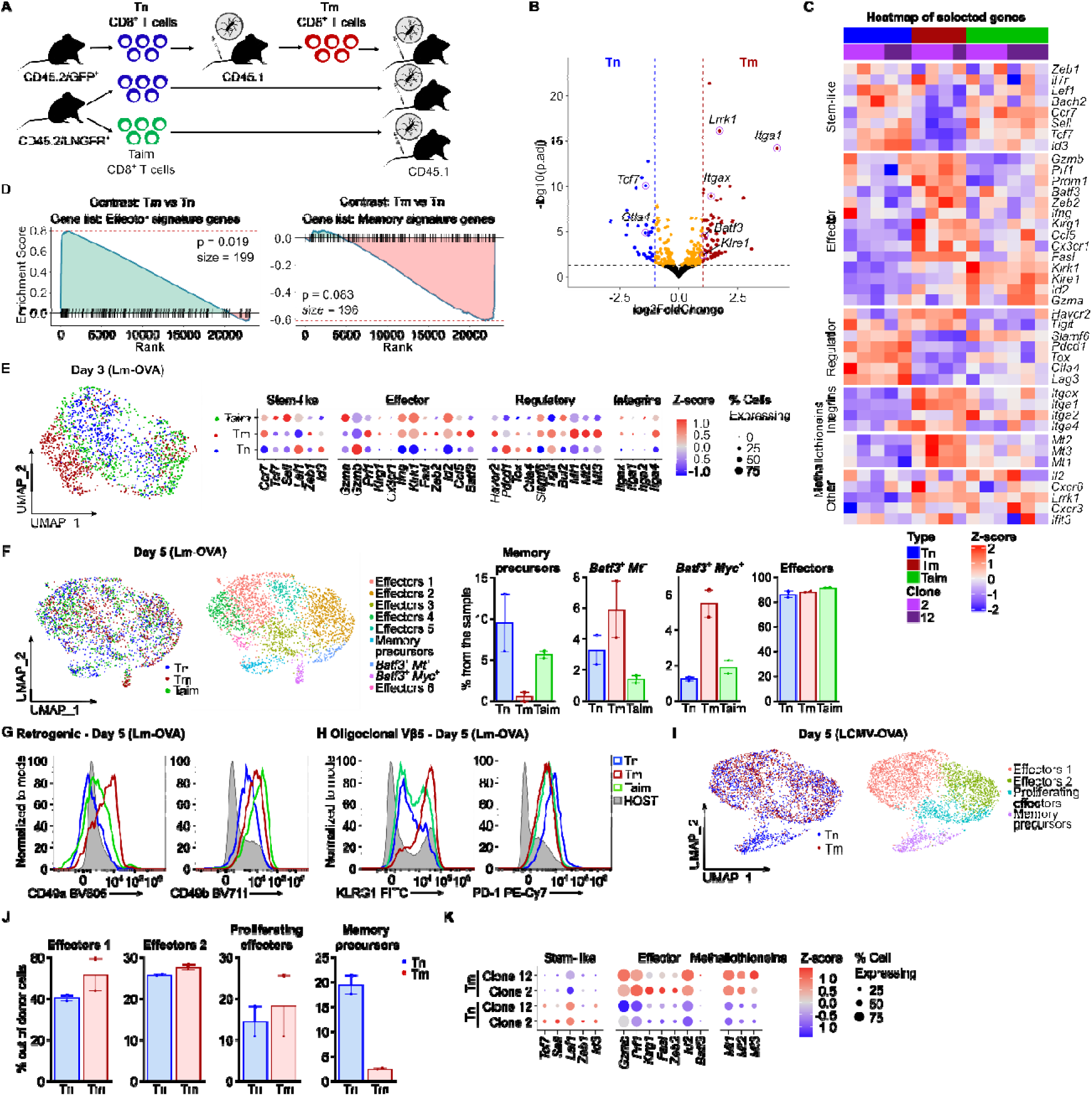
Naive and memory T cells trigger distinct activation programs. **A-G** Sorted Tn (CD8^+^ CD44^-^), Taim (CD8^+^ CD44^+^), and antigen-experienced Tm (CD8^+^ CD45.2^+^) retrogenic T cells (K^b^-OVA-specific retrogenic clone 2 or 12) were adoptively transferred into separate congenic CD45.1 mice followed by Lm-OVA infection. On day 3 or 5 p.i., splenic progeny cells of the donor cells were isolated and analyzed by bulk (B-D), single-cell (E-F) RNA sequencing, or flow cytometry (G). **B-D** Bulk RNA sequencing performed on day 3 p.i. n = 4-6 mice per group. **B** A volcano plot shows differential gene expression between Tn and Tm progeny. Genes with p-adj < 0.05 and |Log2FC| > 1 are highlighted in red (Tm) or blue (Tn). **C** A heatmap of selected genes. **D** Gene set enrichment analysis (GSEA) of differential gene expression between Tn and Tm progeny using effector and memory signature gene lists (GSE1000002_1582_200_UP, GSE1000002_1582_200_DN). **E** ScRNAseq analyses of sorted Tn, Taim, and Tm progeny (clone 2) on day 3 p.i. UMAP showing the cell origin and dot plot of selected genes 3 days p.i. n = 1 mouse per group. **F** ScRNAseq analyses of sorted Tn, Taim, and Tm progeny (clone 2 and 12) on day 5 p.i. UMAP and frequencies of donor cells in particular clusters in individual mice. n = 2 mice per group. **G** FC analysis of Tn, Taim, and Tm progeny (clone 2) on day 5 p.i. **H** Tn (CD8^+^ CD44^-^), Taim (CD8^+^ CD44^+^) or antigen-experienced Tm (CD8^+^ CD45.2^+^) K^b^-OVA-specific T cells from oligoclonal Vβ5 donors were adoptively transferred into congenic CD45.1 mice followed by Lm-OVA infection. Flow cytometry analysis was performed on day 5 p.i. A representative experiment out of 3 in total. **I-K** Sorted Tn (CD8^+^ CD45.2^+^ GFP^+^ CD44^-^) or antigen-experienced Tm (CD8^+^ CD45.2^+^ GFP^+^) retrogenic T cells (clones 2 or 12) were adoptively transferred into CD45.1 congenic mice followed by LCMV-OVA infection. On day 5 p.i., splenic progeny of the donor cells were isolated and analyzed by scRNAseq. n = 2 mice per group. **I** UMAP showing origin of cells and particular clusters. **J** Frequencies of cells in particular clusters in individual mice. **K** Dot plot showing expression of selected genes in individual samples.

We next used single cells RNA sequencing (scRNAseq) to map the cellular states underlying the transcriptional differences identified by bulk RNA-seq. On day 3, the Tm progeny cells differed from Tn and Taim progeny cells, forming predominantly clusters with high expression of *Batf3* and *Mt1* (Fig. 1E, Supplementary Fig. 1G-H). On day 5, most of the progeny formed effectors, irrespective of the original phenotype (Fig. 1F, Supplementary Fig. 1I-J). However, Tn and Taim formed relatively abundant stem-like memory-precursors, whereas Tm progeny were enriched in two *Batf3*-expressing clusters. These results showed substantial differences in the gene expression in Tm and Tn cells and their progeny cells early after activation.

The upregulation of ITGA1 (CD49a), ITGA2 (CD49b), and KLRG1 in Tm than in Tn progeny cells and PD-1 in Tn than in Tm progeny cells was verified by flow cytometry (FC) using monoclonal retrogenic T cells, K^b^-OVA reactive monoclonal OT-I T cells, and K^b^-OVA reactive T cells from oligoclonal Vβ5 mice expressing transgenic OT-I TCRβ and variable endogenous TCRα TCR (Fig. 1G-H. Supplementary Fig. 1K-N).

Next, we performed a similar experiment using LCMV-OVA as a model viral pathogen with scRNAseq analysis on day 5 p.i (Fig. 1I-K, Supplementary Fig. 1O). Again, we observed that Tn cells form significantly higher percentage of memory precursors.

Overall, these experiments reveal specific transcriptional programs in Tn, Taim, and Tm cells triggered by antigenic activation in vivo in the context of bacterial or viral infection. A major difference between Tn and Tm progeny was the preferential formation of stem-like memory precursors by Tn cells and the formation of *Batf3*-expressing subsets and stronger expression of some effector genes by Tm cells.

### Tn cells preferentially generate stem-like Tcm cell precursors

To minimize the effect of the phenotypic heterogeneity of the Tm cells, we further focused on highly proliferative Tcm subset defined as CX3CR1^-^ CD62L^+^. We adoptively transferred Tn and Tcm progeny of K^b^-OVA-specific OT-I *Rag2*^-/-^ T cells into congenic hosts followed by Lm-OVA infection. Next, we characterized their progeny by scRNAseq and flow cytometry. On day 3 post infection, progeny of Tn and Tcm cells were largely distinct and present in separate clusters (Fig. 2A, Supplementary Fig. 2A-B). Even though cell cycle-specific genes were regressed from the data, the unsupervised clustering identified distinct cell clusters which differed in the gene expression profiles as well as in their cell cycle profiles (Supplementary Fig. 2C). Non-dividing cells in G1 phase were mostly coming from the stem-like clusters and were enriched in Tn progeny cells. Together with greater expansion of Tcm cells (Fig. 2C), these data suggested a faster proliferation of Tcm cells than Tn cells. The gene expression analysis revealed that Tn progeny cells express higher levels of genes encoding memory-associated transcription factors (*Tcf7*, *Lef1*, *Id3*), but also checkpoint inhibitors (*Pdcd1*, *Lag3*), cell death receptor (*Fas*) and a key effector cytokine (*Ifng*), whereas Tcm progeny showed higher levels of some effector markers (*Klrg1*, *Klrk1, Ccl5, Fasl*) (Fig. 2B). The expression of major cytotoxic molecules (*Gzma*, *Gzmb, Prf1*) was not greatly varied. We validated these results by FC, where Tcm showed greater expansion and lower expression of PD-1, TCF7, TOX, and FAS proteins than Tn progeny by FC, whereas the GZMB expression was comparable (Fig. 2C, Supplementary Fig. 2D).

**Fig. 2.**
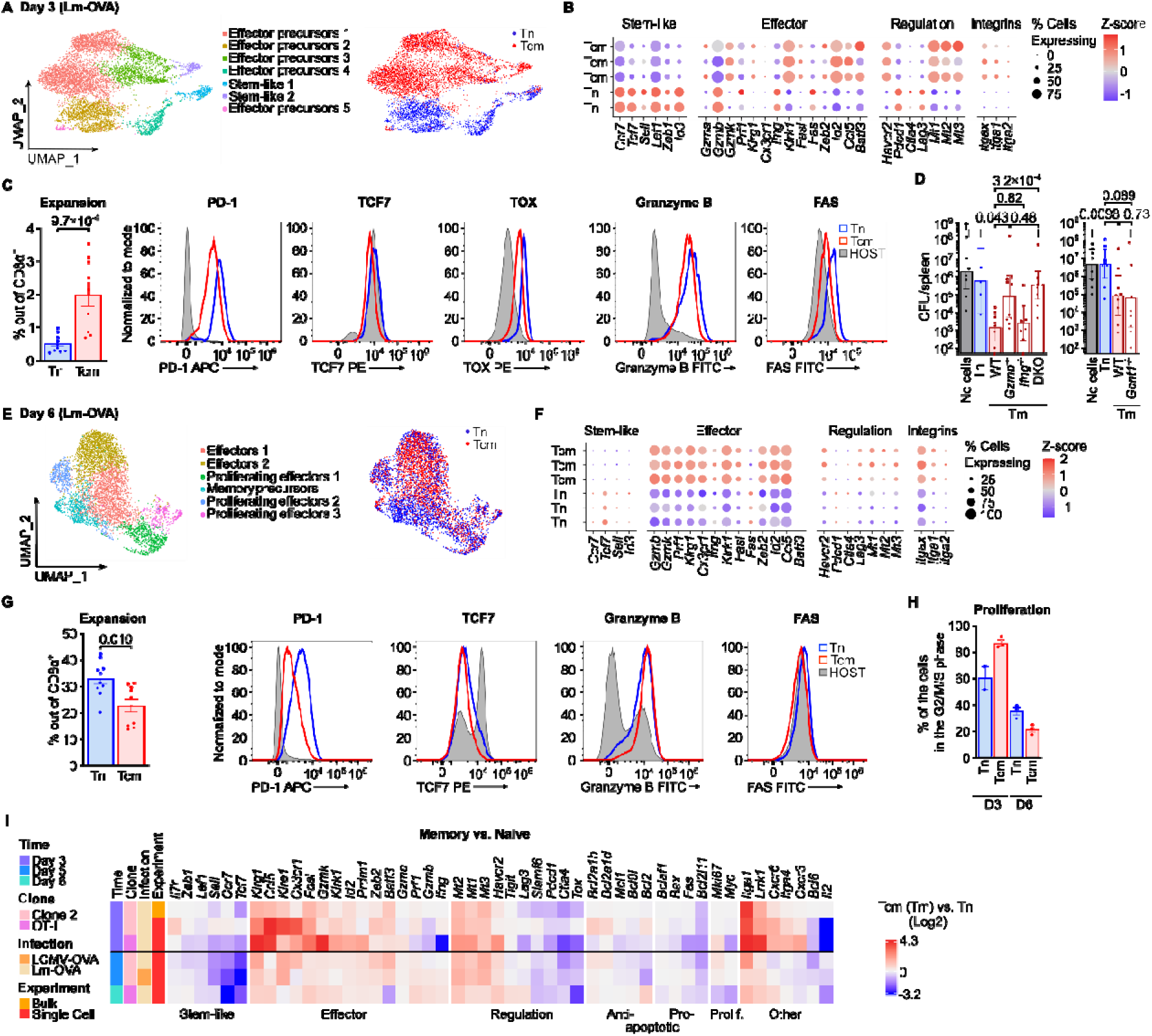
Early activation of Tcm cells drives rapid expansion and effector functions. **A-C** 1.5×10^5^ Tn or Tcm OT-I *Rag2^-/-^* CD8^+^ T cells were adoptively transferred into CD45.1 mice infected with Lm-OVA the next day. Tn and Tcm progeny cells in the spleen were analyzed by scRNAseq (A-B) and FC (C) on day 3 days p.i. **A** UMAP plots showing identified clusters and cell origin. n = 2 (Tn) or 3 (Tcm) mice. **B** Dot plot shows expression of indicated genes in individual samples. **C** Expansion measured as a percentage of donor cells among CD8^+^ T cells (mean ± SEM) and expression of indicated markers. n = 10 (Tn) or 9 (Tcm) in 3 independent experiments. Statistical significance was calculated using two-tailed Mann-Whitney test. **D** 3×10^5^ Tn, or wild-type (WT), *Gzmb^-/-^*, *Ifng^-/-^*, *Gzmb^-/-^ Ifng^-/-^* (DKO), or *Gcnt1^-/-^* Tm OT-I *Rag2^-/-^* CD8^+^ T cells were adoptively transferred to WT mice. Subsequently, mice were infected with high dose of Lm-OVA (1×10^5^ CFU). Splenic bacterial loads were determined on day four p.i. Left: n = 9 (no transfer), 5 (Tn), 8 (WT Tm),10 (*Gzmb^-/-^* Tm), 8 (*Ifng^-/-^* Tm), or 10 (DKO Tm) mice in 3 independent experiments. Right: n = 9 mice in 3 independent experiments. Geometric means with 95 % CI are shown. The dagger symbols (†) represent deceased mice before the measurement. Statistical significance was calculated using two-tailed Mann-Whitney test. Mice with bacterial loads below the detection limit (100 CFU) were given a unique arbitrary value of 50. Deceased animals were given the highest value measured in the same experiment +1. **E-G** 1×10^4^ Tn or 1×10^4^ Tcm OT-I *Rag2^-/-^*CD8^+^ T cells were adoptively transferred into CD45.1 mice followed by Lm-OVA infection. Tn and Tcm progeny cells in the spleens were analyzed by scRNAseq (E-F) and flow cytometry (G) on day six p.i. **E** UMAP plots showing identified clusters and cell origin. n = 3 mice per group. **F** Dot plot shows expression of indicated genes in individual samples. **G** Expansion measured as a percentage of donor cells among CD8^+^ T cells (mean ± SEM) and expression of indicated markers. n = 10 (Tn) or 9 (Tcm) in 3 independent experiments. Statistical significance was calculated using two-tailed Mann-Whitney test. **H** Proportions of cells at G2/M/S cell cycle phases on day 3 and day 6 determined by analysis of scRNAseq data (A and E). Mean ± SEM. **I** A summarizing heatmap of selected genes from all transcriptomic experiments on days 3-6 p.i. showed in Figure 1-2. Relative gene expression is shown as Log2 fold-change between Tcm (or Tm) vs. Tn cells.

In the next step, we focused on key mechanisms of Tm-mediated protection in vivo. In line with previous data ^16^, antigen-specific memory cells provide stronger protection from Lm-OVA infection than Tn cells on per cell basis (Supplementary Fig. 2E). We next used Tm OT-I cells deficient in GCNT1 trafficking factor ^36^, IFNG, GZMB, or both IFNG and GZMB (Fig. 2D, Supplementary Fig. 2E). Whereas we observed that the protection is largely dependent on GZMB, we did not observe any role of GCNT1 and only weak contribution of IFNG on *Gzmb*^+/+^ and *Gzmb*^-/-^ background. The loss of protection by GZMB-deficient Tm cells indicates that early memory-mediated protection was largely dependent on GZMB-mediated cytotoxicity even though the expression of GZMB was not greater in Tm than in Tn cells at this early time point. Plausibly, the initial rapid proliferation of memory progeny contributes to the Lm-OVA clearance, possibly along with high expression of CCL5, which was previously shown to enhance T-cell cytotoxicity ^37^.

On day six p.i, we performed scRNAseq analysis of the progeny of Tn and Tcm cells (Fig. 2E-F, Supplementary Fig. 2F-G). Despite using a conventional method of regressing the cell cycle, we observed multiple clusters differing by the cell cycle phase (Supplementary Fig. 2H). Tcm progeny showed enrichment in the effector clusters, whereas Tn progeny was enriched in the stem-like memory precursors (Supplementary Fig. 2G). In line with the results from day 3, Tn progeny upregulated *Tcf7*, *Pdcd1*, and *Fas*, whereas Tcm progeny upregulated effector markers (*Klrg1, Cxcr3*) and integrins (*Itgax*, *Itga1*) (Fig. 2F-G, Supplementary Fig. 2I). In contrast to day 3, Tcm progeny upregulated cytotoxic genes (*Gzmb, Prf1, Klrk1*) (Fig. 2F-G, Supplementary Fig. 2I). Moreover, the Tn progeny was more abundant than Tcm progeny on day 6 p.i. (Fig. 2G). The quantification of cell cycle showed that Tcm progeny are more proliferative than Tn progeny on day 3 p.i, but this is reversed on day 6 p.i (Fig. 2H).

Overall, we characterized Tn and Tcm responses to cognate infection in vivo at two phases — day 3 and day 5-6 p.i. (Fig. 2I). Three days after infection, Tn cells show modest proliferation and a bias towards stem-like transcriptional responses and Tcm cells undergo rapid proliferation and a bias towards the effector program. Despite relatively weak expression of cytotoxic molecules, Tcm progeny provide protection from cognate intracellular bacteria via direct cytotoxicity on day 4 p.i. This might be explained partially by initially more rapid, but less sustained, proliferation of Tm cells compared to Tn cells. By days 5–6, both populations were dominated by effector states, although Tn-derived cells retained a distinct enrichment of Tcm precursors.

### Tcm-derived progeny persist and preferentially form Tem cells

We examined long-term fates of OT-I Tn and Tcm progeny cells over 12 weeks after Lm-OVA infection. To normalize for the host-to-host variability, we co-transferred congenic CD45.1 Tn and CD45.2 Tcm cells into the same CD45.1/2 heterozygous hosts. Although the peak of expansion in the blood was comparable between Tn and Tcm progeny one-week p.i, we observed a substantial drop of Tn, but not Tcm, progeny at two weeks p.i. (Fig. 3A, Supplementary Fig. 3A-B). The difference in the abundance of Tn- and Tcm-derived progeny persisted throughout the experiment with estimated apparent half-lives of 2.1 and 16.4 days, respectively (Fig. 3A). At one-week p.i, both the Tn and Tcm progeny cells consisted of a majority of Teff/Tem cells gated as CX3CR1^+^ CD62L^-^ or KLRG1^+^ cells. Whereas the percentage of these cells remained high among the Tcm progeny in the following weeks, it substantially dropped in the Tn progeny over the course of the experiment (Fig. 3B). Conversely, the frequency of CX3CR1^-^ CD62L^+^ Tcm cells increased in the Tn progeny more than in the Tcm progeny over time (Fig. 3B). Similar results were observed when Tcm cells were initially sorted as CD62L^+^ independently of CX3CR1 levels (Supplementary Fig. 3C-D).

**Fig. 3.**
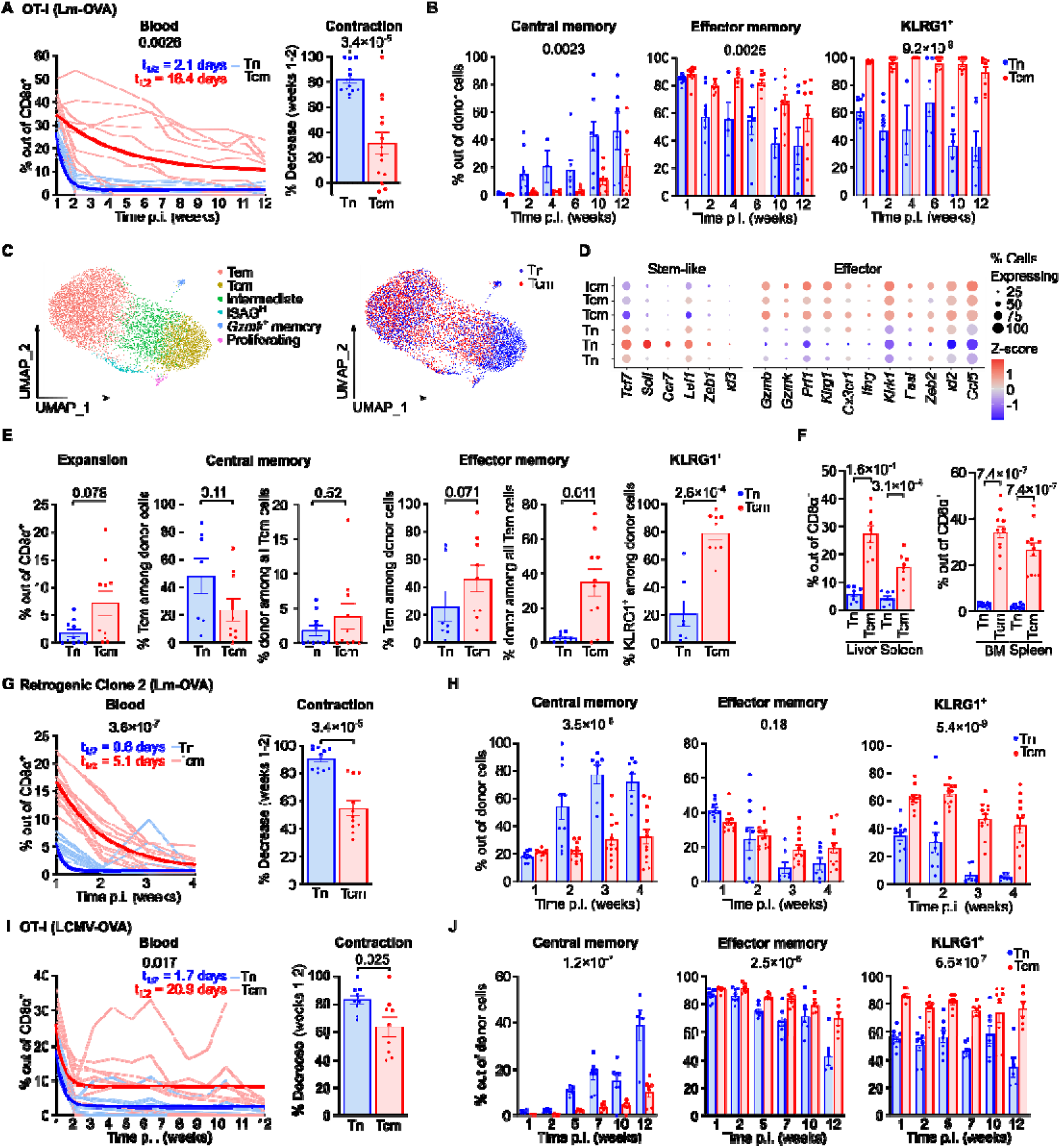
Tn and Tcm cells preferentially generate Tcm and Tem progeny, respectively. **A-E** 1-2×10^4^ Tn (CD8^+^ CD45.1^+^ CD44^-^) and 1-2×10^4^ Tcm (CD8^+^ CD45.2^+^ CD62L^+^ CX3CR1^-^) OT-I *Rag2^-/-^* T cells were co-transferred into congenic hosts (CD45.1/2 heterozygotes) followed by Lm-OVA infection the following day. The cell progeny were analyzed over the course of up to 12 weeks p.i. in the blood and/or spleen by FC and scRNAseq as indicated. **A** The expansion and persistence of Tn and Tcm progeny cells in the blood was determined as their frequency of all CD8α^+^ T cells on a weekly basis. Thin lines show individual donor mice. Thick lines represent fitted nonlinear regression curves (one phase decay) and the half-lives are indicated. n = 10 mice in 2 independent experiments (four mice were sacrificed before the endpoint and are not plotted). The contraction of the cells was calculated as the relative drop in their frequencies between week one and week two p.i. n = 14 mice in 2 independent experiments. **B** Tcm (CD8^+^ CD62L^+^ CX3CR1^-^), Tem (CD8^+^ CD62L^-^ CX3CR1^+^), and KLRG1^+^ cell frequencies among progeny cells in the blood analysis showed in Figure 3A. Only donor progeny subsets comprising of ≥ 0.1 % of CD8^+^ T cells in the blood were included. n = 3-10 mice per group in 2 independent experiments. **C-D** The scRNAseq experiments of Tn and Tcm progeny in the spleen was performed at week 12 p.i. n = 3 per group. **C** UMAP plots showing unsupervised clustering and the cell origin. **D** Dot plot showing the expression of indicated genes in individual samples. **E** Frequencies of Tn and Tcm progeny in all CD8α^+^ T cells and the relative contribution of Tn and Tcm progeny to the overall Tcm, Tem, and KLRG1^+^ frequencies among progeny cells in the host mice were determined by flow cytometry at week 12 p.i. in the spleen. Only donor populations with at least 20 cells measured were included. n = 7-9 mice in 2 independent experiments. **F** 2×10^4^ (liver) or 2.5×10^4^ (BM) of Tn (CD8^+^ CD45.1^+^ CD44^-^) and Tcm (CD8^+^ CD45.2^+^ CD62L^+^ CX3CR1^-^) cells were adoptively co-transferred to congenic hosts subsequently infected with Lm-OVA. The donor progeny cells were detected by FC at 12 weeks (liver) or 32 days (BM) p.i. by FC. Splenic cells from the same donors were analyzed for comparison. n = 8 mice per group in 2 independent experiments (liver) or 12 mice per group in 2 independent experiments (BM). **G-H** 2×10^4^ of Tn (CD8α^+^ CD45.2^+^ LNGFR^+^ CD44^-^) and 2×10^4^ Tcm (CD8α^+^ CD45.2^+^ GFP^+^ CD62L^+^ CX3CR1^-^) retrogenic monoclonal (Clone 2) OVA-specific cells were co-transferred into congenic CD45.1 hosts followed by Lm-OVA infection. Cell progeny were monitored in the blood over the course of 4 weeks p.i. by FC. n = 11-13 mice in 2 independent experiments. **G** Expansion and persistence of the donor progeny. Thin lines show individual donor mice. Thick lines represent fitted nonlinear regression curves (one phase decay). Calculated half-lives are indicated. The contraction of the donor progeny was calculated as the relative drop in their frequencies between week one and week two p.i. **H** The percentage of Tcm, Tem, and KLRG1^+^ cells among the Tn and Tcm progeny was determined by FC at indicated time points. Only donor progeny subsets comprising of ≥ 0.1 % of CD8^+^ T cells in the blood were included. n = 7-13 mice per group in 2 independent experiments. **I-J** 2×10^4^ Tn and Tcm were co-transferred into CD45.1/2 congenic hosts followed by LCMV-OVA infection the following day. Donor progeny were monitored in the blood over the course of 12 weeks p.i. by FC. n = 9 mice in 2 independent experiments. **I** Expansion and persistence of donor progeny cells. Thin lines show individual donor mice. Thick lines represent fitted nonlinear regression curves (one phase decay). Calculated half-lives are indicated. The contraction of the donor progeny cells was calculated as the relative drop in their frequencies between week one and week two p.i. **J** The percentages of Tcm, Tem, and KLRG1^+^ cells among the Tn and Tcm progeny was determined by FC at indicated time points. Only donor progeny subsets comprising of ≥ 0.1 % of CD8^+^ T cells in the blood were included. n = 5-9 mice per group in 2 independent experiments. Statistical significance was calculated using two-tailed Mann-Whitney test for a comparison of two samples (Tn vs Tcm progeny) or two-way ANOVA with mixed-effects analysis (for comparison across multiple time-points).

At the endpoint of 12 weeks p.i., we performed scRNAseq analysis of the Tn and Tcm progeny in the spleen. We identified three major clusters corresponding to Tem (*Cx3cr1*^+^ *Sell*^-^ *Tcf7*^-^), Tcm (*Cx3cr1*^-^ *Sell*^+^ *Tcf7*^+^), and intermediate memory cells (*Cx3cr1*^dim^ Sell^dim^ *Tcf7*^dim^) (Fig. 3C, Supplementary Fig. 3E). Whereas Tem cells were enriched in the Tcm progeny,

Tcm were enriched in the Tn progeny (Fig. 3C, Supplementary Fig. 3F). We also observed three minor memory subsets corresponding to ISAG^hi^ ^35^, proliferating, and *Gzmk*^+^ memory T cells. The latter two were enriched in Tn progeny. Accordingly, the Tn progeny cells showed overexpression of stem-like genes and Tcm progeny overexpressed effector genes (Fig. 3D). FC analysis confirmed the better survival of Tcm progeny and the bias of Tn progeny cells towards Tcm and Tcm progeny cells towards Tem cells (Fig. 3E, Supplementary Fig. 3G). The relative contribution of Tcm and Tn progeny to the Tcm pool was comparable, pointing to the self-renewal ability of Tcm cells during secondary infection (Fig. 3E, Supplementary Fig. 3G). In contrast, the contribution of Tcm progeny to the total Tem pool was dramatically higher than that of Tn progeny. Tcm-derived progeny also outnumbered Tn-derived progeny in the liver (12 weeks) and in the bone marrow (32 days), indicating that their greater persistence was not restricted to blood or secondary lymphoid organs (Fig. 3F).

In agreement with the previous experiments, we observed a rapid contraction as well as biases towards the Tcm phenotype (CD62L^+^ CX3CR1^-^ KLRG1^-^) in Tn progeny and towards the Tem phenotype (CD62L^-^ CX3CR1^+^ KLRG1^+^) in Tcm progeny using retrogenic K^b^-OVA-specific Clone 2 cells responding to Lm-OVA (Fig. 3G-H). Similar differences in the Tn and Tcm progeny survival and fate preferences were observed using OT-I cells and LCMV-OVA infection (Fig. 3I-J, Supplementary Fig. 3H).

Overall, we showed in two monoclonal systems and two infection models that the Tcm-derived progeny persisted more efficiently than Tn-derived progeny and that Tcm progeny is biased towards long-lived Tem formation, whereas Tn progeny is enriched for Tcm cells in the memory phase.

### Tn cell-derived progeny undergo enhanced contraction after peak expansion

Since we observed a rapid reduction of Tn progeny between the first and second week p.i. (Fig. 3A), we monitored the persistence of Tn and Tcm progeny cells on a daily basis (Fig. 4A). We observed a gradual decrease of Tn progeny between days 7 and 14 p.i, which did not occur in Tcm progeny. This was accompanied by an increase in the frequency of CD62L^+^ CX3CR1^-^ Tcm precursors, increase of TCF7^+^ expression and a drop in KLRG1^+^ cells and among Tn progeny (Fig. 4B-C). We did not see a significant difference in CX3CR1^+^ CD62L^-^ Teff/Tem cell frequency between Tn and Tcm donor progeny cells during this time frame. When we infected the mice with LCMV-OVA, the contraction was faster than in the Lm-OVA infection, but the longer persistence of Tcm progeny cells compared to Tn progeny cells was still observed (Supplementary Fig. 4A-B). Because the frequencies of Ki67^+^ cells were comparable between Tn- and Tcm-derived progeny (Supplementary Fig. 4C), we concluded that the differential contraction might be caused by reduced survival of Tn-derived cells.

**Fig. 4.**
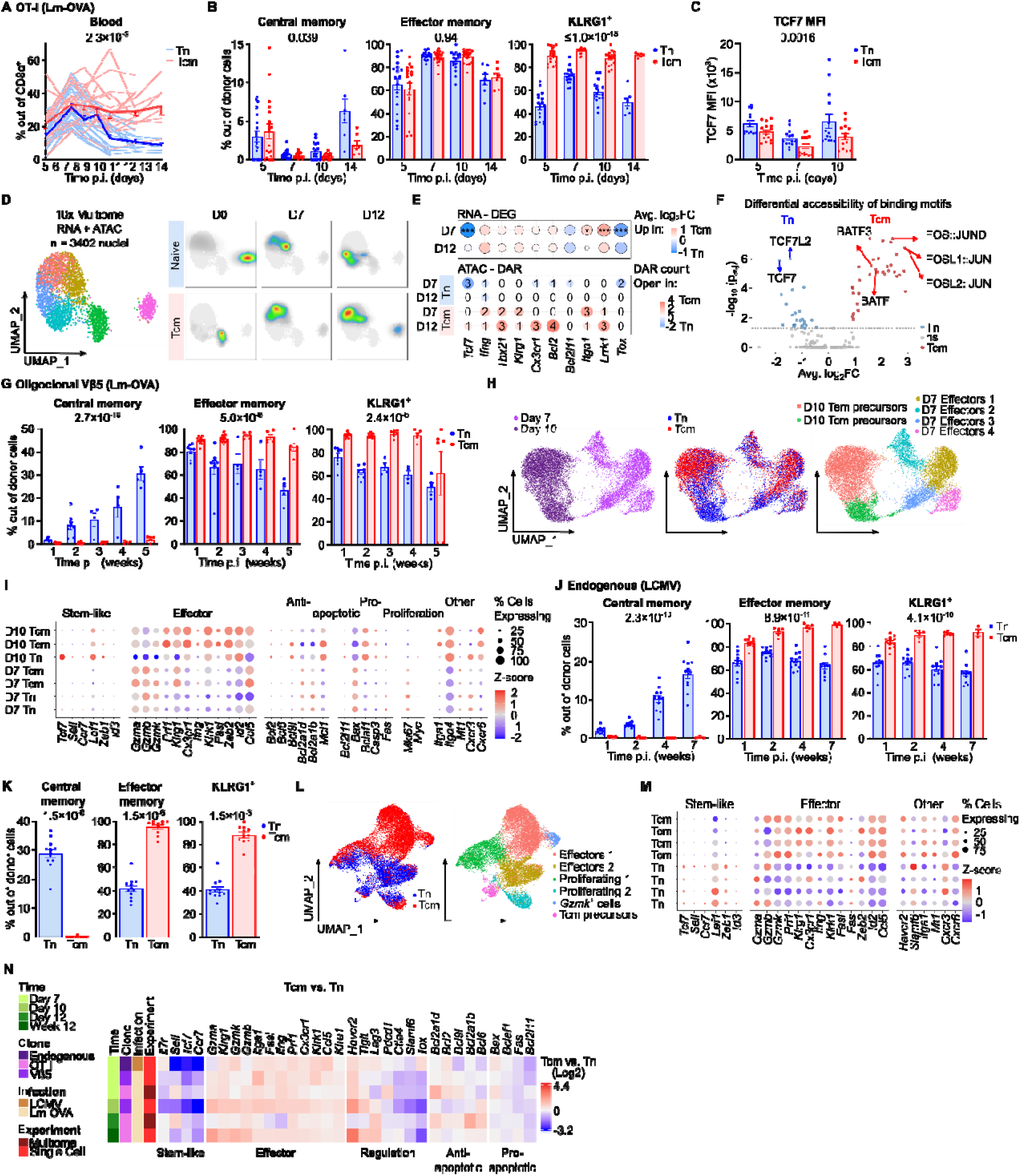
Tn cells consistently upregulate a stemness program across model systems. **A-C** 2×10^4^ sorted Tn (CD8^+^ CD45.1^+^ CD44^-^) and 2×10^4^ Tcm (CD8^+^ CD45.2^+^ CD62L CX3CR1^-^) OT-I *Rag2^-/-^* T cells were co-transferred into congenic host mice (CD45.1/2), followed by Lm-OVA infection. Blood samples were analyzed by flow cytometry on the indicated days. n = 4-18 mice in 4 independent experiments. Data from one of the experiments were included in Figure 3A-B. **A** The expansion and persistence of Tn and Tcm progeny cells in the blood was determined as their frequency of all CD8α^+^ T cells daily. Thin lines show individual donor mice. The thick line represents the mean ± SEM. **B** Tcm (CD8^+^ CD62L^+^ CX3CR1^-^), Tem (CD8^+^ CD62L^-^ CX3CR1^+^), and KLRG1^+^ cell frequences among Tn and Tcm progeny cells in the blood analysis showed in Figure 4A, n = 7-18. **C** Mean fluorescent intensity (MFI) of TCF7 in fixed donor cells. n = 13-14 mice in 3 experiments. **D-F** Nuclei of 1×10^5^ resting Tn and Tm OT-I cells (day 0) or Tn and Tcm OT-I cells responding to Lm-OVA infection isolated on day 7 or 12 p.i. were analyzed by 10x Multiome generating paired scRNAseq and scATACseq data. **D** UMAP based on combined scRNAseq and scATACseq modalities. Colors indicate clusters; pseudocolor density plots show the distribution of the indicated cell populations and time points within the UMAP. **E** The dot plot shows differential expression (DEG) and region accessibility (DAR) of indicated genes. Statistical significance was calculated using Wilcoxon rank-sum test. DEG: * p < 0.05, *** p <0.001. DAR: the count of differentially accessible regions (p < 0.05) within the gene is indicated. **F** The volcano plot shows differential accessibility of transcription factor recognition motifs between Tn and Tcm progeny cells on day 7 p.i. **G-I** 1×10^4^ Tn (CD8^+^ CD45.2^+^ CD44^-^ H-2K^b^-OVA tetramer^+^) or Tcm (CD8^+^ CD45.2^+^ CD62L^+^ CX3CR1^-^) cells from Vβ5 mice were adoptively transferred into separate CD45.1 hosts, followed by Lm-OVA infection next day. Progeny of donor cells in the blood were monitored weekly by flow cytometry (G). On day 7 or 10 p.i., splenic progeny were isolated and analyzed by single-cell RNA sequencing (H, I). **G** Tcm (CD8^+^ CD62L^+^ CX3CR1^-^), Tem (CD8^+^ CD62L^-^ CX3CR1^+^), and KLRG1^+^ cell frequencies among Tn and Tcm progeny cells in the blood. Only the samples with the progeny frequency of at least 0.1 % out of all CD8α^+^cells were considered. n = 4-8 (Tn), 6-13 (Tcm) mice in 3 independent experiments. **H** UMAPs showing the time of isolation of the cells p.i., the cell origin, and identified clusters. Day 7: n = 2 mice per group. Day 10: n = 1 (Tn) or 2 (Tcm). **I** Dot plot showing expression of selected genes in individual samples. **J-M** 1×10^3^ (J-K) or 5×10^3^ (L-M) Tcm (CD8^+^ CD62L^+^ CX3CR1^-^ H-2D^b^-GP33 tetramer^+^) cells were transferred into CD45.1 hosts followed by LCMV infection. Progeny of host H-2D^b^-GP33 tetramer^+^ CD8^+^ T cells and donor Tcm cells were monitored in the blood (J) and spleen (K) by flow cytometry. On day 7 p.i., sorted host and Tcm progeny were analyzed by scRNAseq (L-M). **J-K** Tcm (CD8^+^ CD62L^+^ CX3CR1^-^), Tem (CD8^+^ CD62L^-^ CX3CR1^+^), and KLRG1^+^ cell frequencies among Tn and Tcm progeny cells in the blood (J) and spleen (week 8 p.i.) (K) n = 12 mice per group in 2 independent experiments. Samples with less than 0.1 % Tcm progeny in the blood (n = 6-9) or in the spleen (n = 1) were excluded. **L** UMAPs showing the origin of cells and identified clusters. n = 4 mice. **M** Dot plot shows expression of selected genes in individual samples. **N** Heatmap showing differences in expression of selected genes between Tn and Tcm progeny cells from multiple scRNAseq experiments. If not indicated, a two-way ANOVA with a mixed-effect analysis (comparison of more than 2 samples) or a two-tailed Mann-Whitney nonparametric test were used for statistical analyses. SEM – standard error of the mean

We performed single cell multiome (scRNAseq and scATACseq) analysis of non-activated and activated (day 7 and 12 p.i.) Tn and Tm cells (Fig. 4D). This analysis confirmed increased expression and gene accessibility of stem-like genes such as *Tcf7* in Tn progeny and effector-like genes such as *Klrg1* or *Tbx21* (Tbet) in Tcm progeny (Fig. 4E). Tn-derived progeny showed greater accessibility of TCF7- and TCF7L-family motifs, whereas Tcm-derived progeny showed greater accessibility of AP-1-family motifs, including motifs recognized by JUN/FOS and BATF-family complexes (Fig. 4F). Early activated Tcm progeny cells, but not Tn progeny cells express high levels of an AP-1 family member BATF3 (Fig. 2I), which was previously shown to support survival of memory cells, especially Tem, in the contraction phase ^38^. Since *Batf3* is not expressed in the peak effector phase, it plausibly acts via epigenetic priming during the early activation. Accordingly, we observed a major T-cell pro-apoptotic gene and BATF3-regulated *Bcl2l11* (*Bim*) ^38, 39^ is upregulated and more accessible on day 7 p.i. in the Tn progeny cells in comparison to Tcm progeny cells (Fig. 4E).

### Tn and Tcm cell-derived fate biases are reproduced in oligoclonal and polyclonal responses

In the next step, we compared responses of Tn and Tcm cells outside monoclonal models. First, we transferred K^b^-OVA-specific Tn and Tcm cells isolated from the Vβ5 mice with fixed TCRβ and variable TCRα into congenic host mice, which were subsequently infected with Lm-OVA. In line with the previous data, we observed the biases of Tcm towards CD62L^-^ CX3CR1^+^ KLRG1^+^ Teff/Tem progeny and of Tn towards CD62L^+^ CX3CR1^-^ towards Tcm progeny (Fig. 4G, Supplementary Fig. 4D).

Next, we performed scRNAseq of Tn and Tcm progeny cells on day 7 and 10 p.i. (Fig. 4H, Supplementary Fig. 4E-G). We observed clear differences between these two timepoints reflecting rapid changes in the transcriptome during the contraction phase. Whereas the day 7 p.i. cells contained four clusters of effector cells, which largely differed by cell cycle phases, day 10 p.i. cells separated into Tcm and Tem precursors. The Tcm precursors were enriched in the Tn progeny cells, whereas Tem precursors were enriched in the Tcm progeny cells (Fig. 4H, Supplementary Fig. 4F).

To compare differentiation of Tn and Tcm cells in a fully polyclonal repertoire, we transferred D^b^-GP33(LCMV)-specific Tcm cells into congenic hosts which were subsequently infected with LCMV. We then compared endogenous D^b^-GP33 4mer^+^ T cells and the progeny of transferred Tcm cells by FC and scRNAseq. We considered the endogenous D^b^-GP33 4mer^+^ T cells as predominantly Tn progeny cells, although they possibly contained also progeny of other cell types such as Taim. Moreover, we could not directly compare the magnitude of Tn and Tcm progeny expansion because of their uneven starting numbers and handling (Supplementary Fig. 4H-J). However, we observed again that Tn progeny cells are biased towards Tcm phenotype, whereas Tcm progeny cells are biased towards Teff/Tem phenotype in the blood (Fig. 4J) and the spleen (Fig. 4K).

Our scRNAseq analysis on day 7 p.i. showed a clear distinction between Tn and Tcm progeny (Fig. 4L-M, Supplementary Fig. 4K-M). Tn progeny cells formed Tcm precursors expressing markers of stemness, whereas Tcm progeny cells were biased towards effector-phenotype cells (Fig. 4L-M, Supplementary Fig. 4L-M).

These experiments showed that the differential fate bias of Tn and Tcm progeny was maintained in oligoclonal and polyclonal responses, whereas differences in their long-term persistence were less consistent or could not be directly assessed in these settings.

When all scRNAseq experiments from day 7 to week 12 were put together, the Tn progeny cells consistently showed upregulation of stem-like genes, whereas Tcm progeny upregulated effector genes (Fig. 4N). The expression checkpoint inhibitory receptors also remained different at these later time points with Tn progeny upregulating *Pdcd1* (PD-1), *Ctla4*, *Slamf6* and Tcm progeny upregulating *Havcr2* and *Tigit*. In terms of cell survival, Tn progeny upregulated death receptor *Fas* and pro-apoptotic *Bcl2l11*, whereas Tcm progeny cells upregulated anti-apoptotic *Bcl2* at the peak of expansion (Fig. 4N).

### Tem cells expand poorly but retain a persistent effector phenotype

Beyond the primary comparison of Tn- and Tcm-derived progeny, several experiments included Tem-derived progeny as a separate group or in a Tn/Tem competitive co-transfer setup. Results of these Tem antigenic responses are discussed together in this section.

Some of the above-described experiments included a comparison of Tn and Tem progeny cells. For the sake of simplicity, we discuss their comparison in this separate section chapter.

In the OT-I/Lm-OVA model, Tem cells showed limited early expansion, but also relatively stable numbers over the course of the experiment (Fig. 5A, Supplementary Fig. 5A). ScRNAseq analysis on day 6 p.i. showed that Tem progeny cells lack memory precursors and are enriched for non-proliferating effector cells (Fig. 5B, Supplementary Fig. 5B-D). Accordingly, effector genes were upregulated and memory and stemness genes were down-regulated in Tem progeny cells compared to Tn progeny (Fig. 5C). Tem progeny cells in the blood were almost uniformly CD62L^-^ CX3CR1^+^ KLRG1^+^ from week 1 to week 12 p.i. (Supplementary Fig. 5E-F). At the endpoint of the experiment, Tem progeny cells were highly enriched in the Tem phenotype in the spleen as revealed by scRNAseq and flow cytometry (Fig. 5D-G, Supplementary Fig. 5G). The Tem progeny cells were less abundant than Tn progeny in the liver, which was probably caused by their lower overall expansion (Supplementary Fig. 5H).

**Fig. 5.**
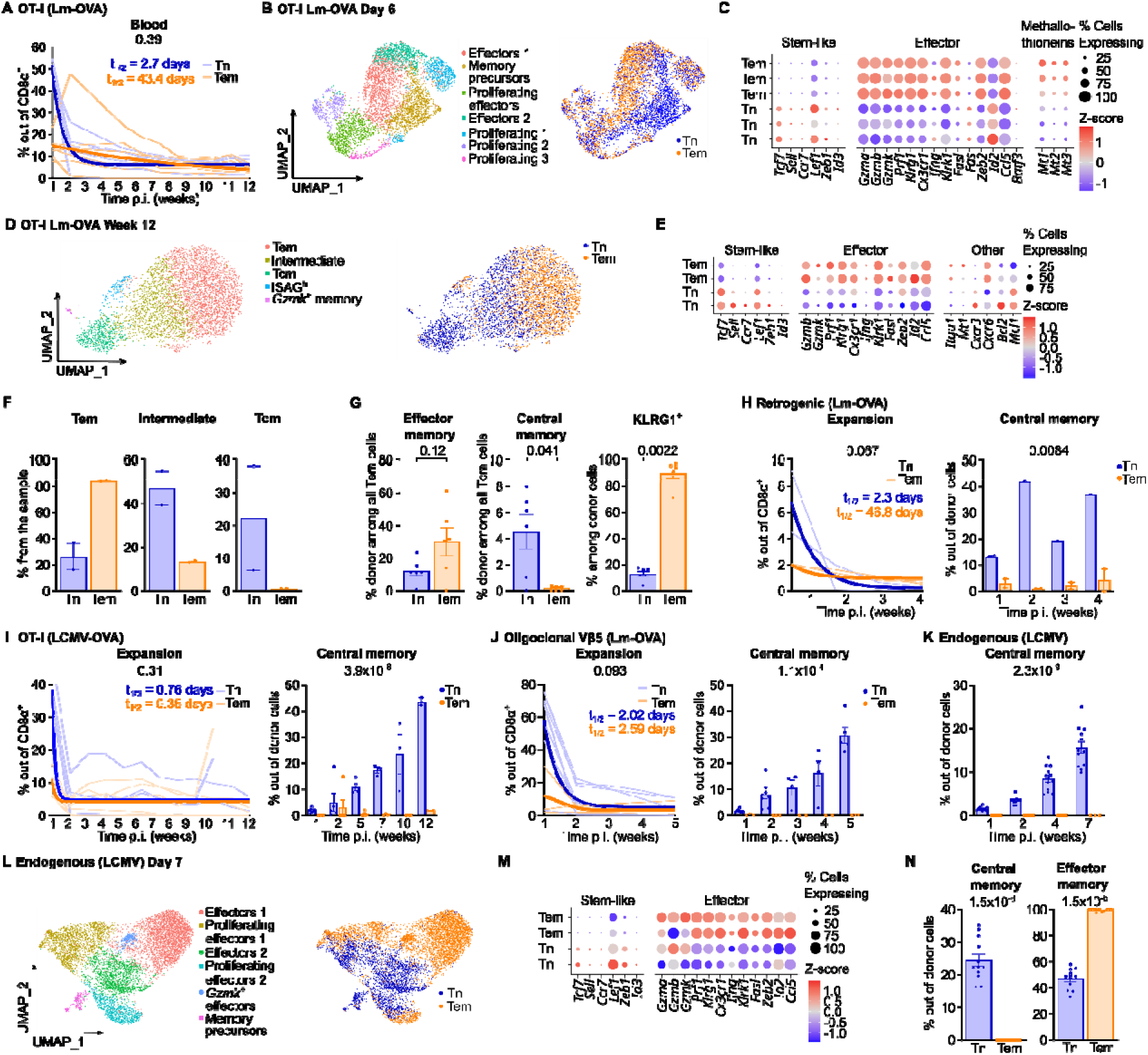
Activated Tem cells show limited proliferation and a restricted fate. **A, D-G** 1-2×10^4^ Tn (CD8^+^ CD45.1^+^ CD44^-^) and 1-2×10^4^ Tem (CD8^+^ CD45.2^+^ CD62L^-^ CX3CR1^+^) OT-I *Rag2^-/-^* cells were co-transferred into congenic CD45.1/2 mice infected with Lm-OVA the following day. The donor progeny cells were analyzed at indicated time points. **A** The expansion and persistence of Tn and Tem progeny cells in the blood was determined as their frequency of all CD8α^+^ T cells on a weekly basis. Thin lines show individual donor mice. Thick lines represent fitted nonlinear regression curves (one-phase decay). The calculated half-lives are indicated. n = 6 mice in 2 independent experiments. **B-C** 1×10^4^ Tn or 1×10^4^ Tem OT-I *Rag2^-/-^*CD8^+^ T cells were adoptively transferred into CD45.1 mice followed by Lm-OVA infection. Tn and Tem progeny in the spleens were analyzed by scRNAseq on day six p.i. Tn samples were used in the analysis shown in Figure 2E-F. **B** UMAP plots showing unsupervised clustering and cell origin. n = 3 mice per group. **C** Dot plot shows expression of indicated genes in the samples. **D** UMAP plots showing unsupervised clustering and cell origin 12 weeks p.i. **E** Expression of indicated genes in the samples 12 weeks p.i. n = 2 from the experiment shown in Figure 3C-D. **F** Frequencies of donor progeny cells in cell clusters12 weeks p.i. **G** The percentage of Tem, Tcm, and KLRG1^+^ cells among the Tn and Tem progeny cells in the spleen at 12 weeks p.i. was determined by FC. n = 6 mice in 2 independent experiments shown also in Figure 5A. **H** 2×10^4^ of Tn (CD8^+^ CD45.2^+^ LNGFR^+^ CD44^-^) and 2×10^4^ Tem (CD8^+^ CD45.2^+^ GFP^+^ CD62L^-^ CX3CR1^+^) retrogenic monoclonal (Clone 2) K^b^-OVA-specific cells were co-transferred into congenic CD45.1 hosts followed by Lm-OVA infection. Cell progeny were monitored in the blood over the course of 4 weeks p.i. by FC. n = 2 mice in 2 independent experiments shown also in Figure 3G-H. Expansion and persistence of the donor progeny. Thin lines show individual donor mice. Thick lines represent fitted nonlinear regression curves (one phase decay). Calculated half-lives are indicated. The percentage of Tcm cells among the Tn and Tem progeny. Samples with less than 0.1 % progeny in the blood (n = 1 Tn) were excluded. **I** 2×10^4^ Tn and Tem were co-transferred into congenic hosts (CD45.1/2 heterozygotes) followed by LCMV-OVA infection the following day. Donor progeny cells were monitored in the blood over the course of 12 weeks p.i. by FC. n = 3-6 mice in 2 independent experiments also shown in Figure 3I-J. Expansion and persistence of the donor progeny. Thin lines show individual donor mice. Thick lines represent fitted nonlinear regression curves (one phase decay). Calculated half-lives are indicated. The percentages of Tcm cells among the Tn and Tem progeny were determined by FC at indicated time points. Only samples with at least 0.1 % progeny in the blood are shown, n = 2-6. **J** 1×10^4^ Tn (CD8^+^ CD45.2^+^ CD44^-^ H-2K^b^-OVA tetramer^+^) or Tem (CD8^+^ CD45.2^+^ CD62L^-^ CX3CR1^+^) Vβ5 CD8^+^ T cells were adoptively transferred into CD45.1 hosts, followed by Lm-OVA infection the following day. Progeny cells in the blood were monitored weekly by flow cytometry. n (Tn) = 8, n (Tem) = 3 mice out of 3 independent experiments also shown in Figure 4G. The percentage of Tcm cells among the Tn and Tem progeny. Samples with less than 0.1 % Tem progeny in the blood (n = 3 Tn) were excluded, or were not measured at weeks 3 and 4 p.i. (n = 4 Tn, 1 Tem). Tn samples were used in analysis shown in figure 4G. **K-N** 1×10^3^ (K, N) or 5×10^3^ (L-M) of Tem (CD8^+^ CD45.2^+^ CD62L^-^ CX3CR1^+^ H-2D^b^-GP33 tetramer^+^) cells were transferred into CD45.1 hosts followed by LCMV infection. Host H-2D^b^-GP33 tetramer^+^ CD8^+^ T cells and donor Tem progeny cells were monitored in the blood (K) and in the spleen at week 8 p.i. (N) by FC. (K, N) n = 12 from 2 independent experiments. Samples with less than 0.1 % Tem progeny in the blood (n = 9) or in the spleen (n = 1) were excluded. (L,N) At day 7 post-infection, sorted progeny from the spleen were analyzed by single-cell RNA sequencing (L-M). n = 2 per group **K** The percentage of Tcm cells among the Tn and Tem progeny cells. **L** UMAP plots showing unsupervised clustering and cell origin. **M** Expression of indicated genes in the individual samples. **N** The percentage of Tem and Tcm cells among the Tn and Tem progeny in the spleen.

The additional experimental setups, i.e., retrogenic Clone2/Lm-OVA (Fig. 5H, Supplementary Fig. 5I), OT-I/LCMV-OVA (Fig. 5I, Supplementary Fig. 5J-L), Kb-OVA^+^ Vβ5/Lm-OVA (Fig. 5J, Supplementary Fig. 5M), and polyclonal D^b^-GP33^+^/LCMV (Fig. 5K-N, Supplementary Fig. 5N-S) consistently showed limited initial expansion and long-term persistence of Tem cells upon cognate infection and strong commitment of their progeny towards the Teff/Tem phenotype, indicating that limited expansion and a persistent Teff/Tem bias were reproducible across several TCR and infection models.

### TCF7 promotes Tcm cell differentiation and TOX regulates PD-1 expression in Tn cell progeny

Our data showed that Tn progeny cells showed a strong intrinsic bias towards formation of Tcm memory precursors during infection. Transcription factors TCF7 and TOX, previously connected to T-cell stemness and longevity in various contexts ^40, 41, 42, 43^, were upregulated in early Tn progeny cells. To evaluate their role in Tn progeny fate trajectories, we knocked-out *Tcf7* or *Tox* genes in Tn OT-I T cells using CRISPR/Cas9 prior to their adoptive transfer to CD45.1 congenic hosts subsequently infected with Lm-OVA. Negative crRNA and Thy1.2-targeting crRNA were used as controls. Knock-out of *Tox* did not alter the frequency of Tcm and Tem cells among the progeny but reduced the expression of PD-1 (Fig. 6A, Supplementary Fig. 6A-B) on day 28 p.i. In contrast, knock-out of *Tcf7* reduced the frequency of Tcm and increased the percentage of Tem progeny cells (Fig. 6A, Supplementary Fig. 6B). These results indicate that TCF7 promotes differentiation of Tn-derived cells toward the Tcm phenotype, whereas TOX contributes to PD-1 upregulation without detectably altering the balance between Tcm and Tem progeny.

**Fig. 6.**
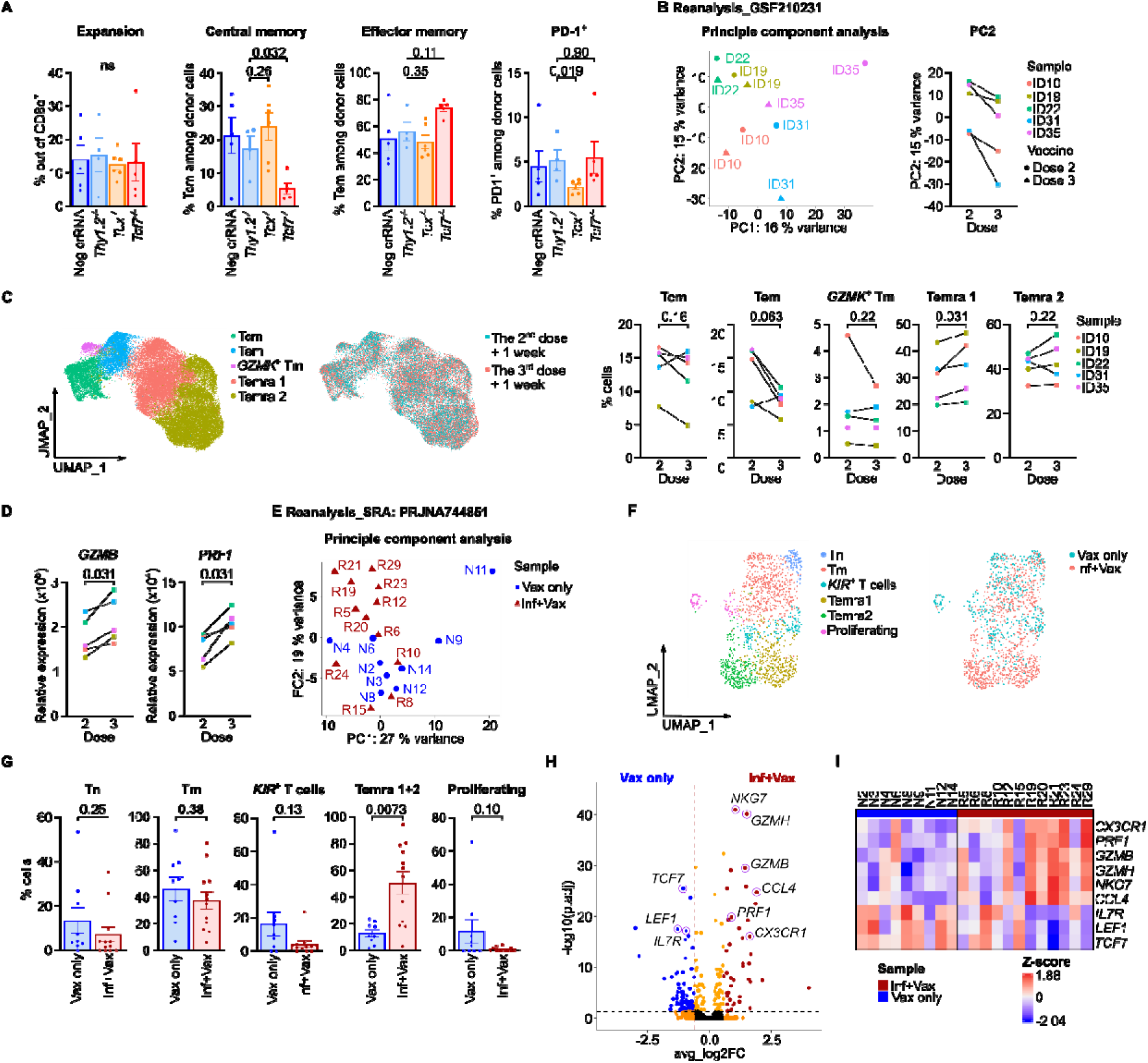
Repeated antigen exposure induces effector programs in human CD8^+^ T cells. **A** Knockout of *Tox*, *Tcf7*, and *Thy1.2* genes (a control together with Neg crRNA – negative control crRNA) was performed on isolated peripheral OT-I T cells using CRISPR/Cas9 technology. 1×10^5^ cells were transferred into congenic CD45.1 mice followed by Lm-OVA infection the day after. The donor progeny cells were analyzed in the spleen 28 days p.i. by FC. Mean and SEM is shown. n = 4-6 per group in 2 independent experiments. Two-tailed Mann-Whitney nonparametric test was used for statistical analyses. SEM – standard error of the mean **B-D** Analysis of human memory T cells from previously published dataset – GSE210231 ^44^. **B** Principal component analysis and difference between the 2nd and the 3rd vaccination dose in PC2, which accounts for 15 % of the variance. **C** UMAP plots showing unsupervised clustering and vaccination dose. Composition of the clusters by individual donors. **D** Relative expression of *GZMB* and *PRF1* expression after the second and third vaccine. **E-I** Analysis of human SARS-CoV-2 spike protein-specific memory CD8^+^ T cells from previously published dataset – SRA: PRJNA744851 ^45^. All donors received 2 doses of the Pfizer-BioNTech BNT162b2 mRNA vaccine. One group of the donors underwent SARS-CoV-2 infection before the vaccination (Inf+Vax), whereas the other group was naive (Vax only). The samples were taken 23-80 days (median 40 days for Inf+Vax and 44 days for Vax only) after the second dose of vaccine. **E** Principal component analysis. **F** UMAPs showing unsupervised cell clustering and donor groups. **G** Relative composition of the clusters. **G** A volcano plot showing differential gene expression between Vax only and Inf+Vax samples. Genes with p-adj < 0.05 and |Log2FC| > Log2(1.5) are highlighted in red (Inf+Vax) or blue (Vax only). **I** A heatmap of selected genes. Statistical significance was evaluated using a one-tailed Wilcoxon paired test (C-D) or two-tailed Mann Whitney test (G).

### Repeated antigen exposure causes an effector-biased shift in human T cells

To examine whether human memory T cells are progressively biased towards the formation of effector/effector memory cells, we reanalyzed a data set with scRNAseq analysis of blood T cells week after the second and third dose of SARS-CoV-2 vaccine ^44^. In the principal component analysis of the gene expression after removing MAIT cells (Supplementary Fig. 6C), the PC1 separated the donors, whereas PC2 highlighted the difference between the second and third dose (Fig. 6B). We identified three major clusters of memory cells and two clusters differentiated effector-like Temra cell (Fig. 6C, Supplementary Fig. 6D). Compared with the response after the second dose, the circulating CD8^+^ T-cell compartment after the third dose showed an increased representation of an effector-like Temra 2 population and higher expression of cytotoxic molecules GZMB and Perforin (Fig. 6C-D). These findings were consistent with an effector-biased shift following repeated vaccination.

To characterize the effect of repeated antigen encounter in antigen-specific cells, we used previously published data ^45^ to compare SARS-CoV-2 spike protein specific CD8+ T cells after two doses of mRNA vaccine in subjects with (Inf+vax) or without (Vax only) a previous SARS-CoV-2 infection event. Despite considerable donor to donor diversity (Fig. 6E), we observed enrichment in the Temra compartment in Inf+Vax donors in comparison to Vax only donors (Fig. 6E-G). Accordingly, we observed upregulation of effector genes GZMB, PRF1, CX3CR1, CCL4 in CD8^+^ T cells in the Inf+vax donors, whereas stemness and memory genes *TCF7*, *LEF1*, and *IL7R* were enriched in Vax only donors (Fig. 6H-I).

## DISCUSSION

Here, we performed a longitudinal single-cell analysis of gene expression changes in CD8^+^ Tn, Tcm, and Tem cells during acute infection across multiple experimental models. Our results show that these CD8^+^ T cells subsets engage distinct cell-intrinsic activation programs during primary and recall responses to cognate antigen. Tn cells preferentially engaged TOX- and TCF7-dependent programs that promoted the early formation of stem-like Tcm precursors and, subsequently, a high frequency of Tcm among their progeny. This finding is consistent with the identification of TCF7^+^ stem-like cells as Tcm precursors throughout the primary CD8^+^ T-cell response ^46^. This program was accompanied by initially slow proliferation, followed by accelerated expansion that generated large numbers of Teff cells by the peak of the immune response. The delayed proliferation may reflect early formation of slow cycling Tcm precursor cells.

Tm cells responded to cognate infection with pronounced differentiation into Teff cells. A related transition was observed in human Tcm and stem cell memory T cells undergoing IL-7- and IL-15-driven proliferation, which induced phenotypic conversion toward Tem cells and increased methylation at the *TCF7* and *CCR7* loci ^19^. Activated Tcm cells triggered rapid initial proliferation, which may facilitate their potent immune protection and is consistent with TCF7-dependent chromatin priming of the metabolic and proliferative programs required during Tcm recall responses ^47^. At the peak of the immune response, Tcm-derived progeny reached numbers comparable to Tn-derived progeny and exceeded those of poorly expanding Tem-derived progeny. However, progeny derived from both Tcm and Tem cells were longer-lived than Tn-derived progeny, resulting in relatively stable populations of Tem cells.

The enhanced persistence of Tcm-derived progeny was most evident in the monoclonal models. A smaller difference was observed in the oligoclonal response, although it did not reach statistical significance because of substantial variability. Persistence could not be directly evaluated in the fully polyclonal model, in which transferred Tcm cells were compared with endogenous antigen-specific cells that differed in precursor abundance, clonal composition, and experimental handling. Nevertheless, the preferential formation of Teff/Tem progeny from Tcm cells and of Tcm progeny from endogenous cells was preserved in this setting. Overall, the progeny-fate bias appears broadly reproducible, whereas the long-term persistence may depend on the clonal and competitive context of the response.

Self-renewal of Tcm cells was compatible with a strong population-level bias toward Teff differentiation. Although Tcm cells can both self-renew and generate differentiated progeny ^11^, only a minority of Tcm-derived cells retained the Tcm phenotype, whereas Tn cells generated Tcm progeny more efficiently. The intermediate activation program engaged by Taim cells, which more closely resembled that of Tn than Tcm cells, further indicates that these differences reflect prior cognate antigen experience rather than the memory-like phenotype alone.

Our experiments in mice and previously published human data ^44, 45^ suggest that repeated antigen exposure may progressively enrich the Teff/Tem compartment while preserving a pool of self-renewing Tcm cells. A similar pattern was observed in a single hypervaccinated individual who had received over 200 SARS-CoV-2 vaccine doses. The frequency of SARS-CoV-2-specific Tcm cells was comparable to that in conventionally vaccinated controls, whereas the frequency of SARS-CoV-2-specific Tem cells was markedly increased ^48^. The ability of repeatedly stimulated T cells to retain proliferative competence after 51 successive immunizations ^49^ further demonstrates that repeated acute stimulations do not deplete the memory pool.

The enhanced survival of Tcm-derived progeny during the contraction phase was associated with higher early expression of *Batf3*. BATF3 was previously shown to promote the survival and memory transition of activated CD8^+^ T cells during contraction by regulating the pro-apoptotic factor BIM, encoded by *Bcl2l11* ^38^. Accordingly, Tn-derived progeny showed greater accessibility at the *Bcl2l11* locus and higher expression of *Bcl2l11* and *Fas*, together with lower expression of the anti-apoptotic gene *Bcl2*, at the onset of contraction.

The balance between Tcm self-renewal and effector differentiation may be altered during sustained antigen exposure, such as in chronic infection or cancer. CAR T cells derived from antigen-experienced Tm cells exhibited superior immediate effector function but reduced proliferative capacity and greater susceptibility to dysfunction than those derived from Tn cells ^50^. Similarly, Tcm-derived progeny failed to sustain proliferation and accumulated to lower numbers than Tn-derived progeny during chronic LCMV infection ^15^. Methallothionein encoding genes *Mt1*, *Mt2*, and *Mt3* were rapidly upregulated in activated Tcm cells. These genes were previously shown to promote exhaustion/dysfunction in tumor infiltrating T cells ^51^. These findings suggest that the differentiation bias towards effector T cells may lead to exhaustion in the conditions of chronic antigen persistence.

Recent studies showed that PD-1^+^ TCF7^+^ TOX^+^ stem-like precursors arise early during both acute and chronic viral infections in naive hosts and can adapt their differentiation to subsequent antigen clearance or persistence ^52, 53^. In cancer, high-avidity stem-like CD8^+^ T cells were recently identified in tumor-draining lymph nodes as TCF7^+^ PD-1^high^ SLAMF6^high^ cells, maintained by PD-1-mediated attenuation of TCR signaling ^54^. The corresponding genes, *Tcf7*, *Pdcd1*, and *Slamf6*, were more highly expressed in early Tn-derived progeny than in Tcm-derived progeny in our study. Although this marker overlap does not establish that the populations are equivalent, it suggests that the bias of activated Tn cells toward stem-like progeny may be particularly important for establishing populations capable of adapting to sustained antigen exposure.

Together, our findings support a model in which Tn and Tm cells trigger different activation programs upon antigen exposure. Thus, prior antigen experience does not merely accelerate a conserved activation response but redirects the cell-intrinsic program that shapes progeny fate and persistence. These distinct programs may reflect adaptation to different histories of antigen exposure. A primary response must preserve long-term regenerative capacity despite uncertain or delayed re-exposure, whereas a recall response occurs to an antigen that has already recurred and may therefore favor rapid effector differentiation and sustained Tem-mediated protection. These findings may inform the design of vaccines and cellular immunotherapies.

## MATERIALS AND METHODS

### Mice

Mice were 6-12 weeks old at the beginning of the experiments. Sex- and age-matched littermates were used, when possible, with females preferred for memory production and males for protection assays.

Mice had a C57BL/6J background, except for *Gzmb^-/-^* (Gzmb^em1(IMPC)Ccpcz^) and *Gcnt1^-/-^* (Gcnt1^em1(IMPC)Ccpcz^) mice, generated on the C57Bl/6NCrl background by the Czech Center for Phenogenomics, Institute of Molecular Genetics of the Czech Academy of Sciences. We further crossed them to the OT-I *Rag2^-/-^* strain on the C57BL/6J background.

IFNγ^OFF^ ^55^, OT I *Rag2^−/−^* ^56, 57^, Vβ5 ^58^, and Ly5.1 ^59^ strains were described previously.

Mice were bred in a specific pathogen-free facility (Institute of Molecular Genetics of the Czech Academy of Sciences; IMG) with a 12-h light/12-h dark cycle and maintained at 22 ± 1 °C and 55 ± 5 % of relative humidity. Mice were fed an irradiated standard rodent breeding diet and quenched with osmosis-filtered water ad libitum. Animal protocols used were in accordance with the laws of the Czech Republic and approved by the Resort Professional Commission for Approval of Projects of Experiments on Animals of the Czech Academy of Sciences, Czech Republic (72/2017 AVCR, 15/2019 AVCR, AVCR 1667/2022 SOV II, AVCR 5262/2024 SOV II).

### Flow cytometry

Organs were harvested and homogenized into cell suspensions. Splenocytes were depleted of erythrocytes by incubation in ACK lysis buffer at RT for 2.5 min. Part of the organ suspension was taken for antibody staining. Cell suspensions and staining were performed in PBS with 2 % FBS, IMDM medium supplemented with 10% FBS (GIBCO) and antibiotics (100 U/ml penicillin (BB Pharma), 100 mg/ml streptomycin (Sigma-Aldrich), 40 mg/ml gentamicin (Sandoz)), RPMI medium supplemented with 10% FBS and antibiotics, or FACS buffer (2 % FBS, 2 mM EDTA, 0.1% sodium azide in PBS) on ice in the dark for 30 minutes.

Prior to intracellular staining, cells were incubated with antibodies against cell-surface markers and fixed using the FOXP3 transcription factor staining buffer set (eBioscience, #00-5523-00).

Blood was collected from the submandibular (facial) vein, depleted of erythrocytes by incubation in ACK lysis buffer at RT for 5 minutes, and intended for antibody staining.

Prior to liver harvest, spleen was isolated and perfusion with PBS through the left heart ventricle was performed. Livers were processed using mouse Liver Dissociation Kit (Miltenyi Biotec, #130-105-807) in gentleMACS Dissociators (Miltenyi Biotec).

Bone marrow was segregated from the femur and tibia by crunching in PBS with 2 % FBS and subjected to ACK erythrocyte lysis for 2 minutes.

### Cells were labelled with antibodies from Biolegend (diluted 1:200)

CD8α (Clone 53-6.7) – BV421 (#100753), AF594 (#100758), PE (#100708), PerCP-Cy5.5 (#100733), PE-Cy7 (#100722); CD45.1 (Clone A20) – APC (#110714), FITC (#110706), BV650 (#110735), AF700 (#110724), PerCP-Cy5.5 (#110728); CD45.2 (Clone 104) – PE (#109808), AF700 (#109822), APC (#109814), FITC (#109806), APC-Cy7 (#109824); CX3CR1 (Clone SA011F11) – PE-Cy7 (#149016), BV711 (#149031), BV605 (#149027); CD62L (Clone MEL-14) – FITC (#104406), AF700 (#104426), PE-Cy7 (#104418); CD44 (Clone IM7) – PE (#103008), BV650 (#103049), FITC (#103006), BV785 (#103059); CD45R (Clone RA3-6B2) – BV510 (#103248); PD-1 (CD279, Clone 29F.1A12) – APC (#135209); FAS (CD95, Clone SA367H8) – FITC (#152605); GRANZYME B (Clone GB11) – FITC (#515403); CD4 – PE (Clone H129.19, #130310), FITC (Clone RM4-4, #116004); CD19 – PE (Clone 6D5, #115508), FITC (Clone 1D3, #152404); KLRG1 (Clone 2F1/KLRG1) – BV510 (#138421), APC (#138412), BV785 (#138429); CD127 (Clone A7R34) – BV421 (#135027); CD49d (Clone R1-2) – PE (#103608), APC (#103622); CD271 (NGFR, Clone ME20.4) – APC (#345108), PE (#345106), Ki-67 (Clone 11F6) – AF647 (#151206).

### Antibodies purchased elsewhere (diluted 1:200)

TOX (Clone TXRX10) – PE (eBioscience, #12-6502-82); TCF7 (Clone S33-966) – PE (BD Pharmingen, #564217); CD45.1 (Clone A20) – PE (BD Pharmingen, #553776); CD45.2 (Clone 104) – PE (BD Pharmingen, #12045483), FITC (BD Pharmingen, #553772); CD49d (Clone R1-2) – PE (BD Pharmingen, #553157).

### Viability staining

LIVE/DEAD™ Fixable Near-IR Dead Cell Stain Kit (Invitrogen, #L34976) was used for the viability staining (1:500).

### Tetramers binding

Tetramers were used for the detection of antigen-specific CD8^+^ T cells. Tetramers were produced by refolding biotinylated monomers and Streptavidin conjugated with PE (Invitrogen, #S866) in a molar ratio of 1:4. Streptavidin–PE was added in two doses with a 30-min incubation on ice after each step. The following biotinylated monomers were used: H-2K^b^-OVA (SIINFEKL) and H-2D^b^-GP33 (KAVYNFATC) from NIH Tetramer Core Facility.

Samples were measured on Cytek Aurora flow cytometer (Cytek) and analyzed using Flow Jo (version 10.6.2, BD Biosciences).

### FACS sorting

Memory CD8^+^ T cells were isolated from the spleens and lymph nodes of previously (≥30 days prior) infected mice. Organs were processed into cell suspensions. Cells were pooled and enriched for CD8^+^ T cells. Fluorescently labeled Tcm (NIR^-^ CD8a^+^ CD45.1^-^ CD45.2^+^ CD62L^+^ CX3CR1^-^) or Tem (NIR^-^ CD8a^+^ CD45.1^-^ CD45.2^+^ CD62L^-^ CX3CR1^+^) and an identical number of naive CD8^+^ T cells (NIR^-^ CD8a^+^ CD44^-^) were FACS sorted and adoptively transferred into congenic hosts.

Antigen-specific oligoclonal and polyclonal CD8^+^ T cells were FACS-sorted using indicated tetramers of biotinylated monomers with Strep-Tactin conjugated with PE (iba, #6-5000-001) or APC (iba, #6-5010-001), which were released by incubating in 1 mM biotin (Sigma, #B4501) solution.

Cell sorting was performed using Aria or Influx sorters (BD Biosciences).

### Cell enrichment

Enrichment of CD8^+^ T cells was performed using the Dynabeads^TM^ Untouched^TM^ Mouse CD8 Cells Kit (Invitrogen, #11417D), with or without homemade rat antibodies anti-B220 and anti-CD4 (YTS177) diluted 1:50.

OT-I T cells from scRNAseq experiment 3 days p.i. were first incubated with antibodies – CD45.1 (Clone A20) – FITC (Biolegend, #110706), CD4 (Clone RM4-5) – biotin (Biolegend, #100508), and CD45R (B220, Clone RA3-6B2) – biotin (Biolegend, #103204) diluted 1:100, then enriched by incubation with anti-FITC magnetic beads (Miltenyi Biotec, #130-048-701) mixed with anti-biotin magnetic beads (Miltenyi Biotec, #130-105-637) for unwanted cells removal by MACS columns.

### RT-qPCR

Total RNA from 10^5^ FACS-sorted cells was isolated using the RNEasy Plus Micro kit (Qiagen, #74034) and transcribed into cDNA using RevertAid reverse transcriptase (Thermofisher, # EP0442) with oligo(dT)18 primers according to the manufacturer’s instructions. cDNA was used for quantitative PCR using the following primers:

SLAMF6 F – tcctttgactagccaacatcc, SLAMF6 R – ggttttgggattttcatttcc;

LAG3 F – ctgagcgatggcagtgtc, LAG3 R – ggtcacctgagattctcctagc;

ITGAX R – caaggggacaacttgagttatga, ITGAX F – tgcttggaatgttcattgtaaaag;

ITGA1 F – ggccaggtcgtcatctaca, ITGA1 R – tgtggttaagacgctaccaaag;

ITGA2 F – tagattccccagcgcaga, ITGA2 R – cgccagacaattcaaaatacc;

BATF F – agcttcagccgctctcct, BATF R – gcagcgatgcgattcttc;

TCF7 F – aggagctgcagccatatgat, TCF7 R – gaggggtttcttgatgactgg;

LRRK1 F – tgtggtggtctggaacctg, LRRK1 R – accagcacaacggcattt;

KLRE1 F – gctccagaccctgctgatta, KLRE1 R – gggttacaggtgcttcatcc;

Eef1a1_F – acacgtagattccggcaagt, Eef1a1_R – aggagccctttcccatctc.

qPCR samples were measured on a LightCycler 480 (Roche) in technical triplicates, and the median value was used for further calculations. The expression of the indicated genes was normalized to the reference gene (Eef1a1) in the same sample.

### Retrogenic T cells

Immortalized bone marrow hematopoietic stem cells from Vβ5 *Rag2^-/-^* mice were transduced with MSCV retroviral vectors encoding Clone 2 or Clone 12 TCRs and GFP or LNGFR selection markers as previously described ^13, 35, 60^. FACS-sorted GFP^+^/LNGFR^+^ CD45.2^+^ cells were transplanted into irradiated (6 Gy) congenic CD45.1 recipient mice using the X-RAD 225XL Biological irradiator (Precision X-Ray). At least 8 weeks after the transplantation, the GFP^+^ or LNGFR^+^ retrogenic T cells were isolated from the spleens and lymph nodes, enriched using Dynabeads^TM^ Untouched^TM^ Mouse CD8 Cells Kit (Invitrogen, #11417D) with in-house made antibodies (anti-B220, anti-CD4 (YTS77; 1:50)), FACS-sorted, and used for subsequent experiments.

### Memory T cells generation

1×10^3^-1.5×10^5^ FACS-sorted naïve (CD8^+^, CD44^-^) cells were adoptively transferred into the tail vein of congenic mice (CD45.1 or CD45.1/2 heterozygotes) followed by Lm-OVA infection. Transplanted cells were of the matching or female sex. At least 30 days post-infection, the cells were resorted from the spleens and lymph nodes as the whole memory pool (Tm, CD8^+^ CD45.2^+^); Tcm (CD62L^+^); Tcm (CD8^+^ CD45.2^+^ (CD44^+^) CD62L^+^ CX3CR1^-^) or Tem (CD8^+^ CD45.2^+^ (CD44^+^) CD62L^-^ CX3CR1^+^) memory and used in downstream analyses.

### Gene knockout in peripheral T cells

Target genes were knockout using Alt-R™ S.p. Cas9-GFP V3 (#1081059), Alt-R CRISPR-Cas9 tracrRNA (#1072533) with two specific crRNAs for each gene: Tox – Mm.Cas9.TOX.1.AA: AACCGGATTCTACCTCATTC (#513636167) and Mm.Cas9.TOX.1.AB: GATCACGGTGTCCAACATGC (#513636166), Tcf7 – Mm.Cas9.TCF7.1.AB: CTGCTGAAATGTTCGTAGAG (#237954613) and Mm.Cas9.TCF7.1.AF: TCTGCTCATGCCCTACCCAC (#237954614), Thy1.2 – Mm.Cas9.THY1.1.AA: CGTGTGCTCGGGTATCCCAA (#512206780) and Mm.Cas9.THY1.1.AC: CCGCCATGAGAATAACACCA (#512206781) or a crRNA negative control (Neg crRNA; #1072544) purchased from Integrated DNA Technologies.

The RNP complexes were assembled by following the manufacturer’s instructions. 1 µl of 60 pmol Cas9-GFP protein was added to each sample (1.8 µl) and incubated for 20 min at RT.

2×10^6^ cells isolated from lymph nodes and spleens of OT-I *Rag2^-/-^* mice were nucleofected using 20 µl of electroporation buffer from the Amaxa^TM^ P4 Primary Cell 4D-Nucleofector™ X Kit S (Lonza, #V4XP-4032) with electroporation enhancer (Integrated DNA Technologies, #1075916) using the DS 137 program. Post nucleofection, cells were cultured under standard conditions (5% CO_2_ at 37 °C) for 1 hour in IMDM medium supplemented with 10% FBS (GIBCO), 100 U/ml penicillin (BB Pharma), 100 mg/ml streptomycin (Sigma-Aldrich), and 40 mg/ml gentamicin (Sandoz).

Afterward, 1×10^5^ cells from each condition were transferred to congenic hosts, followed by Lm-OVA infection. At day 28 p.i., spleens were harvested, processed into cell suspensions, stained with fluorescently labeled antibodies, and measured by flow cytometry.

Cells were counted using the LUNA-II^TM^ Automated Cell Counter (Logos Biosystems).

### Infections

For *Listeria* infection experiments, 5 × 10^3^ CFUs Lm-OVA ^61^ was injected intravenously (i.v.) into the tail vein the day after adoptive cell transfer. In the protection assays, mice were challenged with 1 × 10^5^ CFUs Lm-OVA.

For Lymphocytic choriomeningitis virus (LCMV) infection experiments, mice were injected intraperitoneally with 2×10^5^ PFUs of LCMV Armstrong or LCMV-OVA ^62^ virus. These strains were obtained from Prof. Daniel Pinschewer (University Hospital Basel, Switzerland).

Viral stocks were propagated in BHK-21 cells as described in ^63^. The viral titer of the stocks was determined by LCMV Focus Forming Assay. Briefly, 3T3 cells were infected with different dilutions of the virus. Viral antigens were detected with rat anti-LCMV nucleoprotein antibody (Clone VL-4; BioXCell, #BE0106, diluted 1:500) and visualized by a color reaction using secondary goat anti-rat IgG Horseradish peroxidase antibody (Jackson, #112-035-003, diluted 1:500) 48 h p.i.

### Protection assay

Equal numbers (3×10^5^) of FACS-sorted CD8^+^ OT-I Rag2^-/-^ naive T cells (Tn; NIR^-^ CD4^-^ CD19^-^ CD44^-^ CD45.2^+^), enriched memory T cells (Tm WT; NIR^-^ CD4^-^ CD19^-^ CD45.1^-^ CD45.2^+^), or memory T cells deficient in the production of the target molecules from the spleens were transferred into separate C57Bl/6J mice. Mice were challenged with a high dose of Lm-OVA a day later and sacrificed at day 4 p.i. Spleens were harvested and homogenized in PBS containing 0.1% Tergitol (Sigma, #NP40S). 10-fold serial dilutions of the splenic lysates were plated on brain–heart infusion agar (Sigma, #70138) plates with 200 μg of streptomycin (Carl ROTH, #HP66.1) and incubated at 37°C. Quantification of the bacterial colonies was performed on the following day.

### Bulk RNA sequencing

Memory (CD8^+^ GFP^+^ CD45.2^+^), naive (CD8^+^ LNGFR^+^ CD45.2^+^ CD44^-^), or AIMT (CD8^+^ LNGFR^+^ CD45.2^+^ CD44^+^) retrogenic (Clone 2 and 12) T cells were sorted and transferred into CD45.1 host mice followed with Lm*-*OVA infection. On day 3 post-infection, 2.6×10^4^-5×10^5^ cells were FACS-sorted and the RNA was isolated using Trizol (Invitrogen, #10296028) in combination with the RNEasy plus Micro kit (Quiagen, #74034) according to the manufacturer’s instructions. The cDNA libraries were prepared using the SMARTer Stranded Total RNA-Seq Kit v2 – Pico Input Mammalian (Takara, #634418) according to the manufacturer’s instructions. Single-end sequencing was performed on an Illumina NextSeq 500 using the NextSeq 500/550 High Output Reagent Cartridge v2 (75 cycles) (#20024906), yielding reads of 76 bp. Base calling and FASTQ generation were performed using the Illumina BaseSpace GenerateFASTQ workflow v1.37.0.

### Processing bulk RNA-seq data

Reads were trimmed using TrimmomaticSE ^64^ with the following parameters: ILLUMINACLIP:NexteraPE-PE.fa:2:30:10, LEADING:3, TRAILING:3, SLIDINGWINDOW:4:15, and MINLEN:36; all other parameters were left at their default values. Reads were aligned to the murine reference genome from Ensembl release 102 ^65^ using STAR v2.7.9a ^66^ with default parameters. The reference was prepared according to the STAR user manual using default settings. Reads mapping to the human NGFR sequence (reference generated from GRCh38, Ensembl release 102) were identified using Bowtie2 v2.4.5 ^67^ and removed from the resulting BAM files, as they originated from the reporter construct. Genes encoding TCR and BCR variable segments were removed. Downstream analyses were performed in R v4.5.2 using DESeq2 v1.48.1 ^68^. The PCA plots and heatmaps were generated using design-aware approach. The shrinkage of log2-fold change values was performed using ashr method ^69^. Gene set enrichment analysis (GSEA) was performed using fgsea v1.22.0 ^70^ on gene sets obtained from MSigDB, comparing effector and memory CD8^+^ T cells ^71, 72, 73, 74^. Heatmaps were generated using the ComplexHeatmap package ^75^.

### Single cell RNA-seq experiments

All cells were labeled with one or more hashtag oligonucleotides obtained from BioLegend (TotalSeq™-C). The hashtags used were Mouse Hashtag 1 (Cat. No. 155861), Mouse Hashtag 2 (Cat. No. 155863), Mouse Hashtag 3 (Cat. No. 155865), Mouse Hashtag 4 (Cat. No. 155867), Mouse Hashtag 5 (Cat. No. 155869), Mouse Hashtag 6 (Cat. No. 155871), Mouse Hashtag 7 (Cat. No. 155873), Mouse Hashtag 8 (Cat. No. 155875), Mouse Hashtag 9 (Cat. No. 155877), Mouse Hashtag 10 (Cat. No. 155879), Mouse Hashtag 11 (Cat. No. 155881), and Mouse Hashtag 12 (Cat. No. 155883). Cells from all experiments were loaded onto a Chromium or Chromium X Controller (10x Genomics). The targeted cell recovery per sample depended on the experiment. cDNA libraries for the first two experiments were prepared using the Chromium Single Cell V(D)J Reagent Kits with Feature Barcode technology for Cell Surface Protein protocol (#CG000186 Rev D) and the Chromium Single Cell 5′ Library & Gel Bead Kit and Chromium Single Cell 5′ Feature Barcode Library Kit (10x Genomics; #PN-1000014, #PN-1000020, #PN-1000080, #PN-1000009, #PN-1000084).

The remaining experiments, except for the final one, were prepared using the Chromium Next GEM Single Cell 5′ Reagent Kits v2 (Dual Index) with Feature Barcode technology for CRISPR Screening and Cell Surface Protein (10x Genomics; #PN-1000263, #PN-1000286, #PN-1000541, #PN-1000190, #PN-1000215, #PN-1000250), following the manufacturer’s protocol (#CG000511 Rev C). For the final experiment, cDNA libraries were prepared using the Chromium GEM-X Single Cell 5′ Reagent Kits v3 with Feature Barcode technology for CRISPR Screening and Cell Surface Protein (10x Genomics; #PN-1000694, #PN-1000698, #PN-1000699, #PN-1000703, #PN-1000215, #PN-1000250), following the manufacturer’s protocol (#CG000736 Rev A). Sequencing was performed on an Illumina NovaSeq 6000 platform.

### Processing and analyzing single cell RNA-seq data

The murine reference genome used for read alignment was obtained from Ensembl release 102 and prepared using the *mkref* tool from 10x Cell Ranger v5.0.1 ^76^. Count matrices were generated using the *count* tool from either 10x Cell Ranger v5.0.1 or v8.0.1, using either R2-only or paired-end sequencing data, depending on the experiment.

Subsequent analyses were performed in R v4.5.2 using the Seurat v5.3.0 package ^77^. Cells not assigned to any expected hashtag combination were removed. Mitochondrial genes, ribosomal genes, genes encoding TCR variable segments, and genes detected in fewer than three cells were excluded. Cells with more than 10% of reads mapping to mitochondrial genes or expressing fewer than 200 genes after gene filtering were removed.

Each dataset was normalized using Seurat’s default normalization method with a scale factor of 1×10, scaled, and subjected to dimensionality reduction by principal component analysis (PCA) and uniform manifold approximation and projection (UMAP), followed by Louvain clustering. Parameters for PCA, UMAP, and clustering were optimized separately for each dataset. Clusters consisting predominantly of dead cells or contaminating cell types were removed, after which the analysis pipeline was repeated from the normalization step onward.

Some preprocessed datasets were subsequently split by experimental day. Samples sequenced across multiple wells were integrated using the STACAS v2.3.0 package ^78^ or, in the case of Week 12 samples, merged directly. Any newly identified contaminating clusters were removed, and the datasets were reprocessed. Cell cycle effects were regressed out using the gene list from the scGate package ^79^, except in the Week 12 samples.

Differential expression analysis was performed using the FindAllMarkers and FindMarkers functions from Seurat, employing a two-sided Wilcoxon rank-sum test with Bonferroni correction for multiple testing. Heatmaps comparing log fold changes between Tcm/Tm and Tn cells were generated using the ComplexHeatmap package.

### Multiome analysis

FACS-sorted Tcm (CD8a^+^ CD45.1^-^ CD45.2^+^ CD62L^+^ CX3CR1^-^), Tem (CD8a^+^ CD45.1^-^ CD45.2^+^ CD62L^-^ CX3CR1^+^), and naïve cells (CD8a^+^ CD44^-^) were adoptively transferred into congenic (CD45.1) host followed by Lm-OVA infection ending up with 2 mice per condition. In indicated data points, spleens were harvested and processed into cell suspension. Donor cells were resorted (CD8a^+^ CD45.1^-^ CD45.2^+^ CD44^+^) together with resting memory (CD8^+^ CD45.2^+^ CD44^+^) and naïve (CD8^+^ CD44^-^) cells. Samples were pooled together based on the condition.

Nuclei were isolated according to the 10x Genomics Demonstrated Protocol CG000365 (Rev C), with minor adaptations. Briefly, samples were washed in PBS containing 0.04% BSA and lysed in chilled lysis buffer for 3 minutes on ice. For sample multiplexing, isolated nuclei from each sample were labelled with MULTI-seq Lipid-Modified Oligos kit (Sigma-Aldrich, #LMO001A, #LMO001B) according to the original protocol ^80^. Sample-specific MULTI-seq barcode oligonucleotides were custom synthesized and used in the format 5′-CCTTGGCACCCGAGAATTCCA-[8-nt sample barcode]-poly(dA)30-3′. The sample barcode sequences used were: GGCTGCGC, TAGTTGAC, GACGCGCG and CGTCCTAG. The resulting nuclei were washed two times in wash buffer, pooled and resuspended in diluted 1× nuclei buffer supplemented with 1 mM DTT (Sigma-Aldrich, #646563) and 1 U/μl RNase inhibitor (Sigma-Aldrich, #3335402001), and counted immediately before loading.

Single-nucleus ATAC and gene expression libraries were generated using the Chromium Next GEM Single Cell Multiome ATAC + Gene Expression reagents (10x Genomics, #PN-1000285) according to the manufacturer’s protocol (CG000338, Rev B), except for modifications introduced to recover the lipid-hashtag barcode library. Transposition, GEM generation, barcoding, post-GEM cleanup, pre-amplification, and ATAC library construction were performed according to the standard workflow. During cDNA amplification, the standard 10x cDNA amplification master mix was supplemented with MULTI-seq primer (CTTGGCACCCGAGAATTCC) to enable amplification of the sample barcode oligonucleotides ^80^. In the subsequent cDNA cleanup step, the 0.6× SPRI supernatant was retained as the lipid-hashtag barcode fraction. The bead-bound endogenous cDNA was processed further according to the standard 10x protocol and used for gene expression library construction. The retained hashtag barcode fraction was transferred to a fresh tube and purified by high-ratio SPRI selection with isopropanol, eluted in EB buffer, and amplified by PCR using TruSeq RPI primers + Universal I5 primer to generate a separate hashtag barcode library. ATAC, gene expression, and hashtag barcode libraries were quantified and quality-controlled using Agilent Bioanalyzer and sequenced on NextSeq 2000 using NextSeq 1000/2000 P2 Reagents (100 cycles) kit (Illumina, #20046811).

### Processing Multiome data

10x Multiome data were analyzed using Seurat (v5.1.0) and Signac (v1.13.0). Cell Ranger ARC filtered feature-barcode matrices and ATAC fragment files were loaded, mouse genome annotations were added from EnsDb.Mmusculus.v79/mm10 (v2.99.0), and cells were quality-filtered based on RNA counts (nCount_RNA > 100 and < 45000), ATAC counts (nCount_ATAC > 100 and < 100000), TSS enrichment (> 1) and nucleosome signal (< 2.5). Hashtag-derived sample annotations were manually assigned based on hashtag counts and transferred to the multiome object by barcode. Peaks were re-called from ATAC fragments using MACS2, quantified with FeatureMatrix, and stored as a custom peak assay. RNA data were normalized with SCTransform and reduced by PCA, whereas ATAC data were processed by TF–IDF normalization and LSI; joint or accessibility-based UMAPs and clusters were then generated. D7 and D12 objects were merged and hashtag identities representing Tem cells were excluded from the final analysis. Motifs were annotated using JASPAR2020/mm10 (v0.99.10) followed by Signac chromVAR motif-activity scoring. Differential RNA expression, chromatin accessibility and motif activity between Tcm and naïve cells at D7 and D12 were assessed with Seurat FindMarkers; differential peaks were annotated to nearby genes using ClosestFeature/LinkPeaks.

### Processing public human single-cell data sets

All datasets were processed from raw count matrices made available by the authors of the respective publications using R v4.5.2, Seurat v5.3.0, DESeq2 v1.48.1, and STACAS v2.3.0.

For the dataset GEO: GSE210231 ^44^, only samples from the second experimental batch were analyzed. RNA and surface protein (ADT) assays from individual wells were normalized independently (log-normalization for RNA and CLR normalization for ADT), scaled, and subjected to principal component analysis (PCA; 50 and 10 principal components for RNA and ADT, respectively). Batch correction was then performed separately for the RNA and ADT modalities using Seurat’s layer integration framework (CCA integration for RNA and RPCA integration for ADT). The integrated modalities were combined using weighted nearest neighbor (WNN) analysis. UMAP was computed from the weighted nearest-neighbor graph using the default WNN parameters, and cells were clustered using the SLM algorithm with a resolution of 0.2. Cell clusters were manually annotated. MAIT cells were subsequently removed, and the entire analysis was repeated using identical parameters. Differential expression analysis was performed using the Seurat function *FindMarkers* with the Wilcoxon rank-sum test. For pseudobulk analysis, raw counts were aggregated by sample and analyzed using DESeq2. PCA was calculated with using design-aware approach.

For the dataset published by Minervina et al. ^45^, only the cells with spike-specific response, and only from donors taken from either naïve subjects, or recovered donors taken at second sampling (after infection and both vaccine doses) were analyzed. The mitochondrial, ribosomal (including mitochondrial ribosomal) genes, and genes encoding TCR variable segments were removed. The data set was then normalized, scaled, subjected to PCA (20 principal components) and UMAP dimensional reduction and clustering (resolution of 0.2).

The volcano plot was computed from values generated using FindMarkers for contrast comparing vaccinated and recovered donors vs. vaccinated donors, and it includes only genes that were present in at least 10% of either group. The analyses comparing individual subjects and pseudobulk analyses were performed only on donors with at least 20 cells.

## Acknowledgments

We gratefully acknowledge Ladislav Cupak for technical assistance, Zdenek Cimburek and Matyas Sima (Flow cytometry facility, IMG) for cell sorting, Sarka Kocourkova and Michal Kolar (Laboratory of Genomics and Bioinformatics, IMG) for cDNA library preparations.

## Funding

Czech Science Foundation grant 22-18046S (OS)

European Union’s Horizon 2020 research and innovation programme ERC Starting Grant ActSwiftly under grant agreement No. 101125695 (OS)

Czech Health Research Council NW26A-NIVB (OS) Charles University Grant Agency 177024 (ES)

Core funding provided by the Institute of Molecular Genetics of the Czech Academy of Sciences RVO68378050 (OS, RS)

European Union and the Ministry of Education, Youth and Sports of the Czech Republic (MEYS) M2023036 and CZ.02.01.01/00/23_015/0008189 (Upgrade of the large research infrastructure CCP III) (RS)

## Author contributions

Conceptualization: ES, DP, JM, OS

Methodology: ES, DP, JM, VN, AD, MK, RS, OS

Investigation: ES, DP, JM, VN, VC, KT, AD, VR, AM, AMM, OS

Visualization: ES, DP, JM, VN, OS

Funding acquisition: ES, RS, OS

Project administration: OS Supervision: RS, OS

Writing – original draft: ES, JM, OS

Writing – review & editing: ES, DP, JM, VN, VC, KT, AMM

## Data and materials availability

All datasets generated in this study are available from the NCBI Gene Expression Omnibus (GEO) under accession number GSE342096.

The human data reanalyzed in our study are available from NCBI GEO under accession number GSE210231^44^ and at https://zenodo.org/records/6232103 and https://github.com/pogorely/COVID_vax_CD8^45^.

The code used to analyze data generated in this study and the publicly available datasets is available on GitHub at https://github.com/Lab-of-Adaptive-Immunity/MEMOs.

