## Supplementary material for "Distinct activation programs in naive and memory CD8 T cells govern progeny stemness and effector persistence": Fig. S

**Supplemental Material**

**Supplemental Figures 1-6**

### Distinct activation programs in naïve and memory CD8 T cells govern stemness and effector persistence of their progeny

**Eva Salyova[1,2]*, Darina Paprckova[1]*, Juraj Michalik[1], Veronika Niederlova[1], Veronika Cimermanova[1], Katerina Tomicova[1], Ales Drobek[1], Vojtech Racek[1], Alena Moudra[1], Michaela Krupkova[3], Radislav Sedlacek[3], Ondrej Stepanek[1]**

1 Laboratory of Adaptive Immunity, Institute of Molecular Genetics of the Czech Academy of Sciences, Prague, Czechia

2 Faculty of Science, Department of Cell Biology, Charles University, Prague, Czechia

3 Czech Centre for Phenogenomics & Laboratory of Transgenic Models of Diseases, Institute of Molecular Genetics of the Czech Academy of Sciences, Vestec, Czechia

* These authors contributed equally to this study

### Figure S1

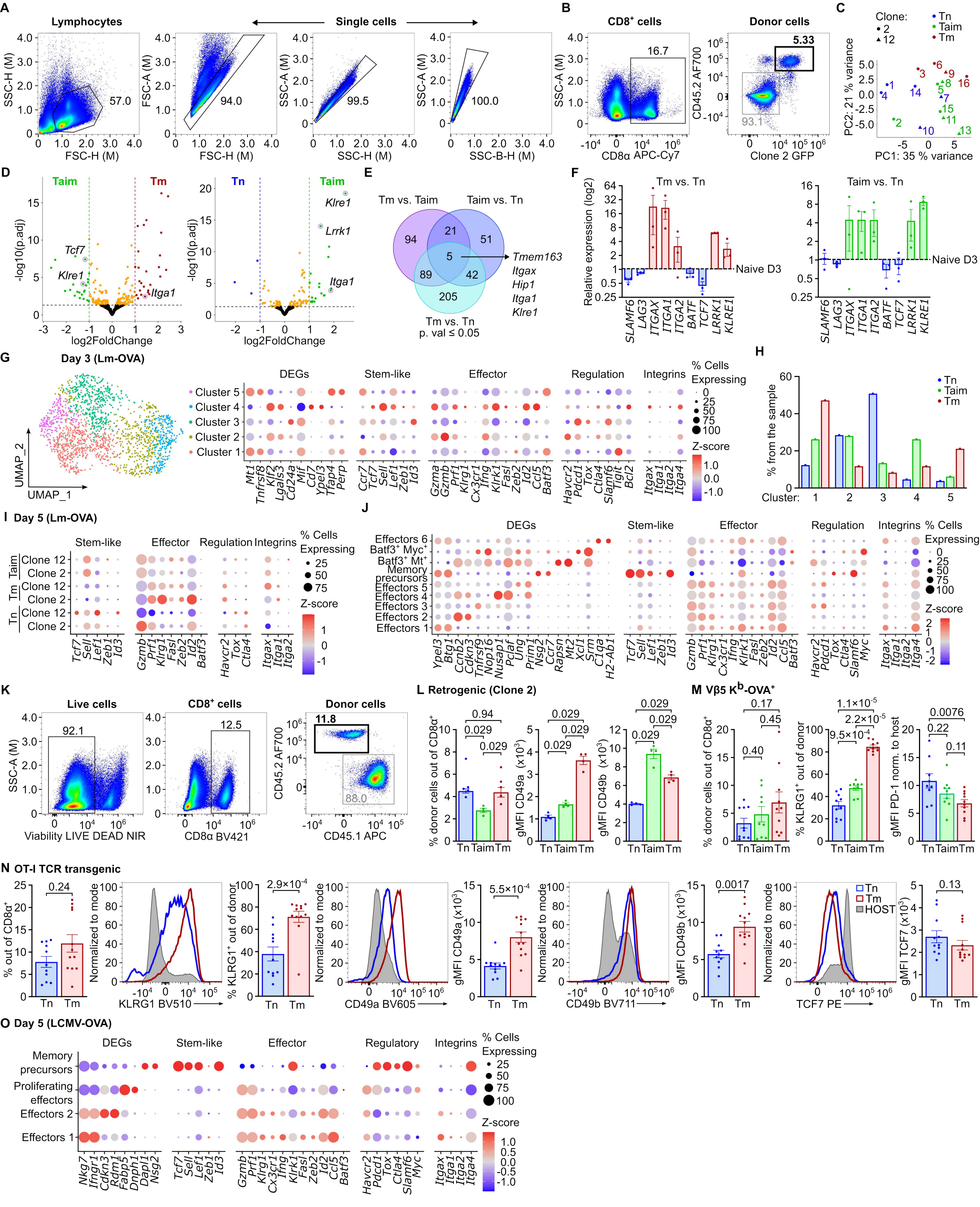

A
Lymphocytes
Single cells
SSC-H (M)
FSC-H (M)
FSC-A (M)
SSC-A (M)
SSC-B-H (M)
57.0
94.0
99.5
100.0
B
CD8+ cells
Donor cells
16.7
5.33
93.1
CD8α APC-Cy7
CD45.2 AF700
Clone 2 GFP
C
Clone:
2
12
Tn
Taim
Tm
PC1: 35 % variance
PC2: 21 % variance
D
Taim
Tm
Tn
Tcf7
Klre1
Itga1
Lrrk1
-log10(p.adj)
log2FoldChange
E
Tm vs. Taim
Taim vs. Tn
Tm vs. Tn
p. val ≤ 0.05
94
21
51
5
89
42
205
Tmem163
Itgax
Hip1
Itga1
Klre1
F
Tm vs. Tn
Taim vs. Tn
Relative expression (log2)
Naive D3
SLAMF6
LAG3
ITGAX
ITGA1
ITGA2
BATF
TCF7
LRRK1
KLRE1
G
Day 3 (Lm-OVA)
UMAP_1
UMAP_2
Cluster 5
Cluster 4
Cluster 3
Cluster 2
Cluster 1
DEGs
Stem-like
Effector
Regulation
Integrins
% Cells Expressing
Z-score
H
% from the sample
Cluster:
I
Day 5 (Lm-OVA)
Clone 12
Clone 2
J
Effectors 6
Batf3+ Myc+
Batf3+ Mt+
Memory precursors
Effectors 5
Effectors 4
Effectors 3
Effectors 2
Effectors 1
K
Live cells
CD8+ cells
Donor cells
92.1
12.5
11.8
88.0
Viability LIVE DEAD NIR
CD8α BV421
CD45.1 APC
L
Retrogenic (Clone 2)
% donor cells out of CD8α+
gMFI CD49a (x10³)
gMFI CD49b (x10³)
0.94
0.029
M
Vβ5 Kb-OVA+
0.17
0.45
0.40
1.1×10⁻⁵
2.2×10⁻⁵
9.5×10⁻⁴
0.0076
0.22
0.11
% KLRG1+ out of donor
gMFI PD-1 norm. to host
N
OT-I TCR transgenic
0.24
2.9×10⁻⁴
5.5×10⁻⁴
0.0017
0.13
% out of CD8α+
Normalized to mode
KLRG1 BV510
CD49a BV605
CD49b BV711
TCF7 PE
HOST
gMFI TCF7 (x10³)
O
Day 5 (LCMV-OVA)
Memory precursors
Proliferating effectors
Effectors 2
Effectors 1
DEGs
Stem-like
Effector
Regulatory
Integrins

**Figure S1**

**A** Representative gating strategy of singlets used in this study.

**B** Representative gating of retrogenic T cells as ($CD45.2^+$, $GFP^+$).

**C-E** Additional analyses of the RNAseq experiment on progenies of retrogenic Tn, Tm, and Taimt cells on day 3 p.i. presented in Figure 1B-D.

**C** Principal component analysis.

**D** Volcano plots showing contrasts between Taim vs. Tm progenies and Tn vs. Taim progenies.

**E** Numbers of differentially expressed genes in the sample comparisons and their intersections.

**F** Relative expression of indicated genes determined by RT-qPCR using the same RNA samples as in C-E. The relative expression in Tm and Taim cells is shown as a fold change to Tn levels. n = 3 (2 – Clone 2, 1 – Clone 12) independent experiments.

**G-H** Additional analyses of the scRNAseq experiment on progenies of retrogenic Tn, Tm, and Taim cells on day 3 p.i. (Lm-OVA) presented in Figure 1E.

**G** UMAP and dot plot of selected genes.

**H** Frequencies of the clusters in individual mice.

**I-J** Additional analyses of the scRNAseq experiment on progenies of retrogenic Tn, Tm, and Taim cells on day 5 p.i. (Lm-OVA) presented in Figure 1F.

**I** A dot plot of indicated genes showing differences among samples.

**J** A dot plot of indicated genes showing differences among clusters.

**K** A gating strategy for gating on adoptively transferred $CD8^+$ T cells from Vβ5 mice and OT-I *Rag2*$^{-/-}$ in experiment shown in Figure 1H, Figure S1M-N.

**L-N** Flow cytometry analysis of expansion and indicated markers on progeny of Tn, Taim, and Tm cells from (L) retrogenic (Clone 2) cells (shown in Figure 1G), n = 4 mice per group, (M) oligoclonal Vβ5 mice (shown in Figure 1H), n = 9-10 mice per group, and (N) monoclonal OT-I *Rag2*$^{-/-}$ mice, n = 11-12 mice per group.

**O** Dot plot of indicated genes showing differences among clusters. An additional analysis of the scRNAseq experiment on progenies of retrogenic Tn and Tm on day 5 p.i. (LCMV-OVA) presented in Figure 1I-K. n = 2 mice per group.

Indicated P-values were caculated by two-tailed Mann–Whitney test. DEGs – differentially expressed genes.

### Figure 2S

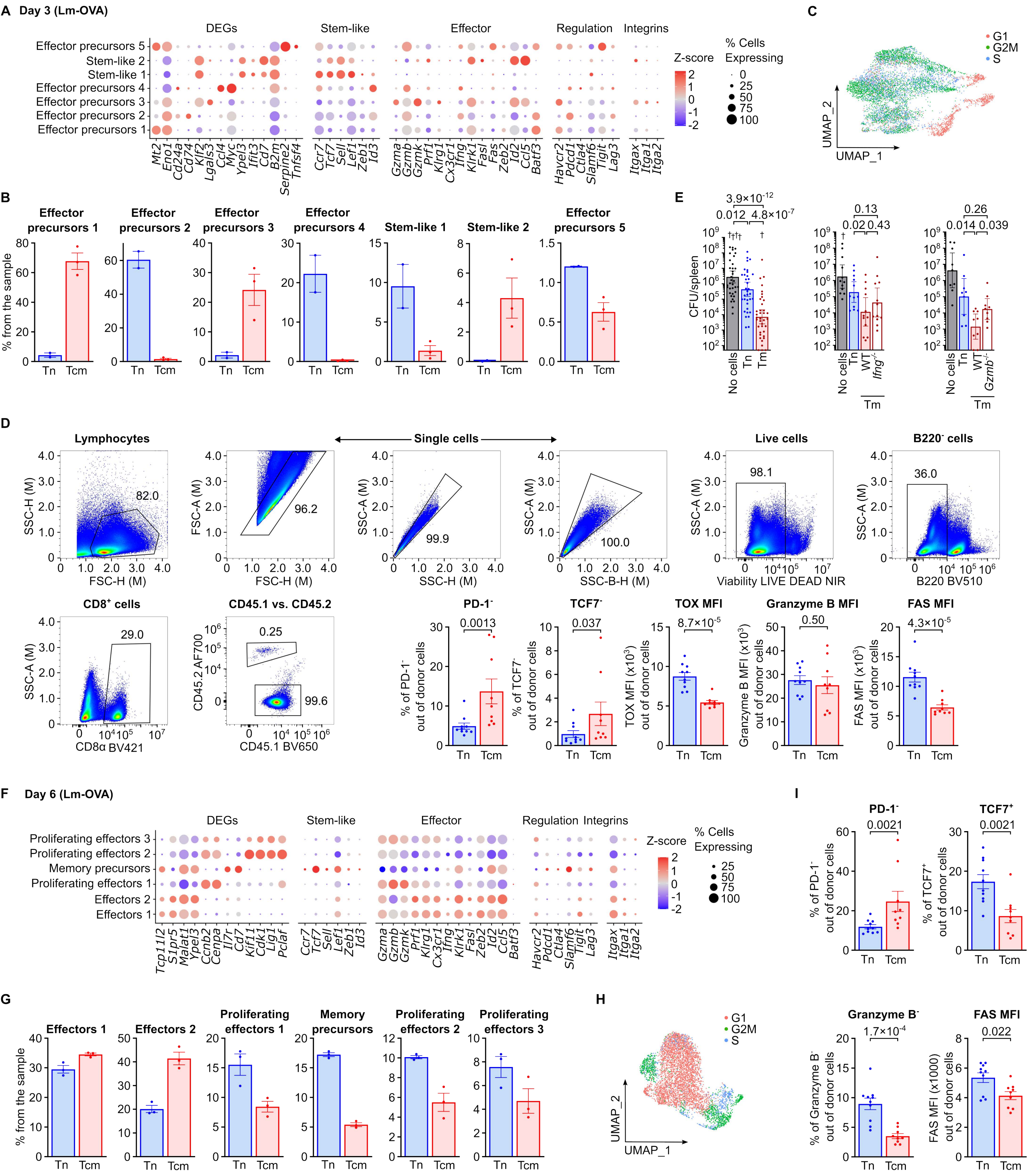

A Day 3 (Lm-OVA)
DEGs
Stem-like
Effector
Regulation
Integrins
Effector precursors 5
Stem-like 2
Stem-like 1
Effector precursors 4
Effector precursors 3
Effector precursors 2
Effector precursors 1
Mt2 Eno1 Cd24a Cd74 Klf2 Lgals3 Ccl4 Myc Ypel3 Ifit3 Cd7 B2m Serpine2 Tnfsf4
Ccr7 Tcf7 Sell Lef1 Zeb1 Id3
Gzma Gzmb Gzmk Prf1 Klrg1 Cx3cr1 Ifng Klrk1 Fas Fasl Zeb2 Id2 Ccl5 Batf3
Havcr2 Pdcd1 Ctla4 Slamf6 Tigit Lag3
Itgax Itga1 Itga2
Z-score
% Cells Expressing
C
G1
G2M
S
UMAP_2
UMAP_1
B
Effector precursors 1
Effector precursors 2
Effector precursors 3
Effector precursors 4
Stem-like 1
Stem-like 2
Effector precursors 5
% from the sample
Tn Tcm
E
3.9×10^-12
0.012
4.8×10^-7
0.13
0.02
0.43
0.26
0.014
0.039
CFU/spleen
No cells Tn Tm
No cells Tn WT Ifng-/-
No cells Tn WT Gzmb-/-
Tm
D
Lymphocytes
Single cells
Live cells
B220- cells
82.0
96.2
99.9
100.0
98.1
36.0
SSC-H (M)
FSC-H (M)
FSC-A (M)
SSC-A (M)
SSC-H (M)
SSC-B-H (M)
Viability LIVE DEAD NIR
B220 BV510
CD8+ cells
CD45.1 vs. CD45.2
29.0
0.25
99.6
CD8α BV421
CD45.2 AF700
CD45.1 BV650
PD-1-
TCF7-
TOX MFI
Granzyme B MFI
FAS MFI
0.0013
0.037
8.7×10^-5
0.50
4.3×10^-5
% of PD-1- out of donor cells
% of TCF7- out of donor cells
TOX MFI (x10^3) out of donor cells
Granzyme B MFI (x10^3) out of donor cells
FAS MFI (x10^3) out of donor cells
F Day 6 (Lm-OVA)
Proliferating effectors 3
Proliferating effectors 2
Memory precursors
Proliferating effectors 1
Effectors 2
Effectors 1
Tcp11l2 S1pr5 Malat1 Ypel3 Ccnb2 Cenpa Il7r Cd7 Kif11 Cdk1 Lig1 Pclaf
Ccr7 Tcf7 Sell Lef1 Zeb1 Id3
Gzma Gzmb Gzmk Prf1 Klrg1 Cx3cr1 Ifng Klrk1 Fasl Zeb2 Id2 Ccl5 Batf3
Havcr2 Pdcd1 Ctla4 Slamf6 Tigit Lag3
Itgax Itga1 Itga2
I
PD-1-
TCF7+
0.0021
0.0021
% of PD-1- out of donor cells
% of TCF7+ out of donor cells
G
Effectors 1
Effectors 2
Proliferating effectors 1
Memory precursors
Proliferating effectors 2
Proliferating effectors 3
H
Granzyme B-
FAS MFI
1.7×10^-4
0.022
% of Granzyme B- out of donor cells
FAS MFI (x1000) out of donor cells

**Figure 2S**

**A** Expression of indicated genes in cluster (data from Fig. 2A).

**B** Frequencies of cells in indicated clusters composition by the samples (data from Fig. 2A).

**C** UMAP shows cell cycle stages (data from Fig. 2A).

**D** Representative gating strategy for the flow cytometry experiments in Fig. 2. Quantification of indicated markers in experiment presented in Fig. 2C shown as frequencies of negative cells or mean fluorescent intensities (MFI). n = 10 (Tn) or 9 (Tcm) mice in 3 independent experiments. Two-tailed Mann-Whitney test was used for the calculation of the statistical significance.

**E** The protection assay was performed as described in Fig. 2D. The first plot aggregate data from all performed experiments. n = 39 (no transfer), 35 (Tn), or 38 (TM) mice in 13 experiments. The following plots show results not overlapping with data shown in Fig. 2D.

GCNT1: n = 9 mice per group. IFNG: n = 12 mice per group. GZMB: n = 9 mice per group.

Geometric means with 95 % CI are shown. Statistical significance was calculated using two-tailed Mann-Whitney test.

**F** Expression of indicated genes in cluster (data from Fig. 2E).

**G** Frequencies of cells in indicated clusters composition by the samples (data from Fig. 2E).

**H** UMAP shows cell cycle stages (data from Fig. 2E).

**I** Quantification of indicated markers in experiment presented in Fig. 2G shown as frequencies of positive and negative cells or mean fluorescent intensities (MFI). n = 10 (Tn) or 9 (Tcm) mice in 3 independent experiments. Two-tailed Mann-Whitney test was used for the calculation of the statistical significance.

**Figure 3S**

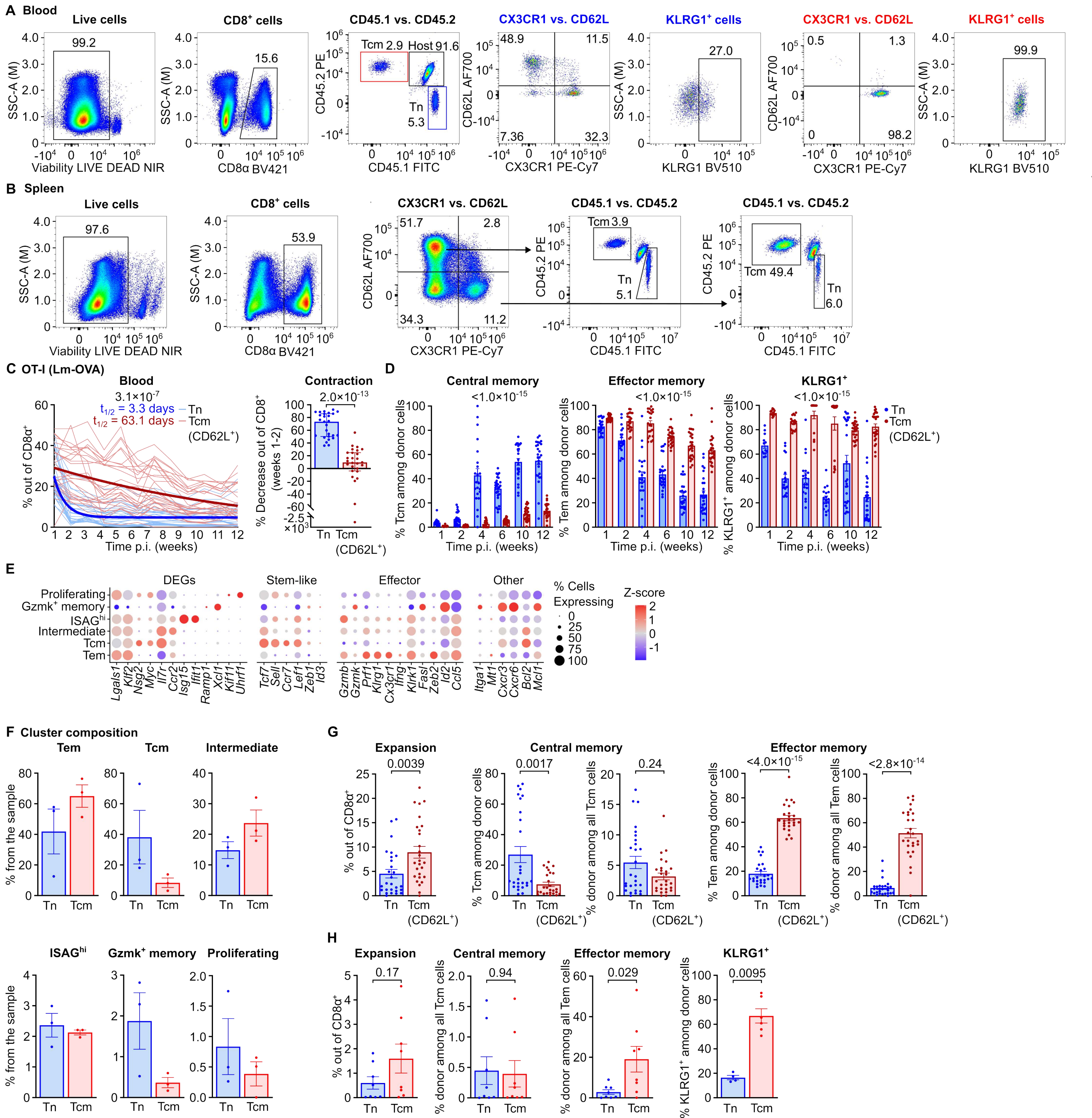
A Blood
Live cells
CD8+ cells
CD45.1 vs. CD45.2
CX3CR1 vs. CD62L
KLRG1+ cells
B Spleen
C OT-I (Lm-OVA)
Blood
Contraction
D
Central memory
Effector memory
KLRG1+
E
DEGs
Stem-like
Effector
Other
% Cells Expressing
Z-score
F Cluster composition
Tem
Tcm
Intermediate
ISAGhi
Gzmk+ memory
Proliferating
G
Expansion
Central memory
Effector memory
H
Expansion
Central memory
Effector memory
KLRG1+

**Figure S3**

**A-B** Representative gating strategies for determining the persistence and phenotype of blood (A) and splenic (B) Tn and Tcm progeny cells. A strategy capturing intrinsic differentiation of donor cells (A) and a strategy showing the contribution of donor cells to the Tcm and Tem compartments (B) are shown.

**C-D, G** $2.5x10^4$ Tcm ($CD8α^+$ $CD62L^+$ $CD44^+$ $CD45.2^+$) cells were co-transferred with $2.5x10^4$ of Tn ($CD8α^+$ $CD44^-$ $CD45.1^+$) cells into CD45.1/2 hosts, infected with Lm-OVA the following day. The donor progeny cells were analyzed in the blood (C-D) and spleen (G) by FC.

**C** The expansion and persistence of Tn and Tcm progeny in the blood was determined as their frequency of all $CD8α^+$ T cells on a weekly basis. Thin lines show individual donor mice. Thick lines represent nonlinear regression curves (one phase decay) and the calculated half-lives are indicated. The contraction of the donor progeny cells was calculated as the relative drop in their frequencies between week one and week two p.i. Median with interquartile range is shown. n = 27 mice in 3 independent experiments.

**D** Tcm ($CD8^+$ $CD62L^+$ $CX3CR1^-$), Tem ($CD8^+$ $CD62L^-$ $CX3CR1^+$), and $KLRG1^+$ cell frequencies among donor progeny cells in the blood analysis showed in Figure S3C. Only the samples with the progeny frequency of at least 0.1 % out of all $CD8α^+$ cells were considered. n = 13-26 mice in 2 independent experiments.

**E** Expression of indicated genes in cell clusters in the scRNAseq experiment shown in Figure 3C.

**F** Frequencies of donor progeny cells in cell clusters for each individual donor population in the scRNAseq experiment shown in Figure 3C.

**G** The analysis of donor progeny cells in the spleen by FC at 12 weeks p.i. n = 27 mice in 3 independent experiments. Only donor populations with at least 20 cells were included, n = 26 mice.

**H** The percentage of Tcm, Tem, and $KLRG1^+$ cells among the Tn and Tcm progeny in the spleen at 12 weeks post LCMV-OVA infection was determined by FC at indicated time points. n = 8 mice in the 2 experiments shown in Figure 3I-J. For the quantification of $KLRG1^+$ cells, only donor populations with at least 20 cells measured were included, n = 4-6.

Statistical significance was calculated using two-tailed Mann-Whitney test for a comparison of two samples (Tn vs Tcm progeny) or two-way ANOVA with mixed-effects analysis (for comparison across multiple time-points).

### Figure 4S

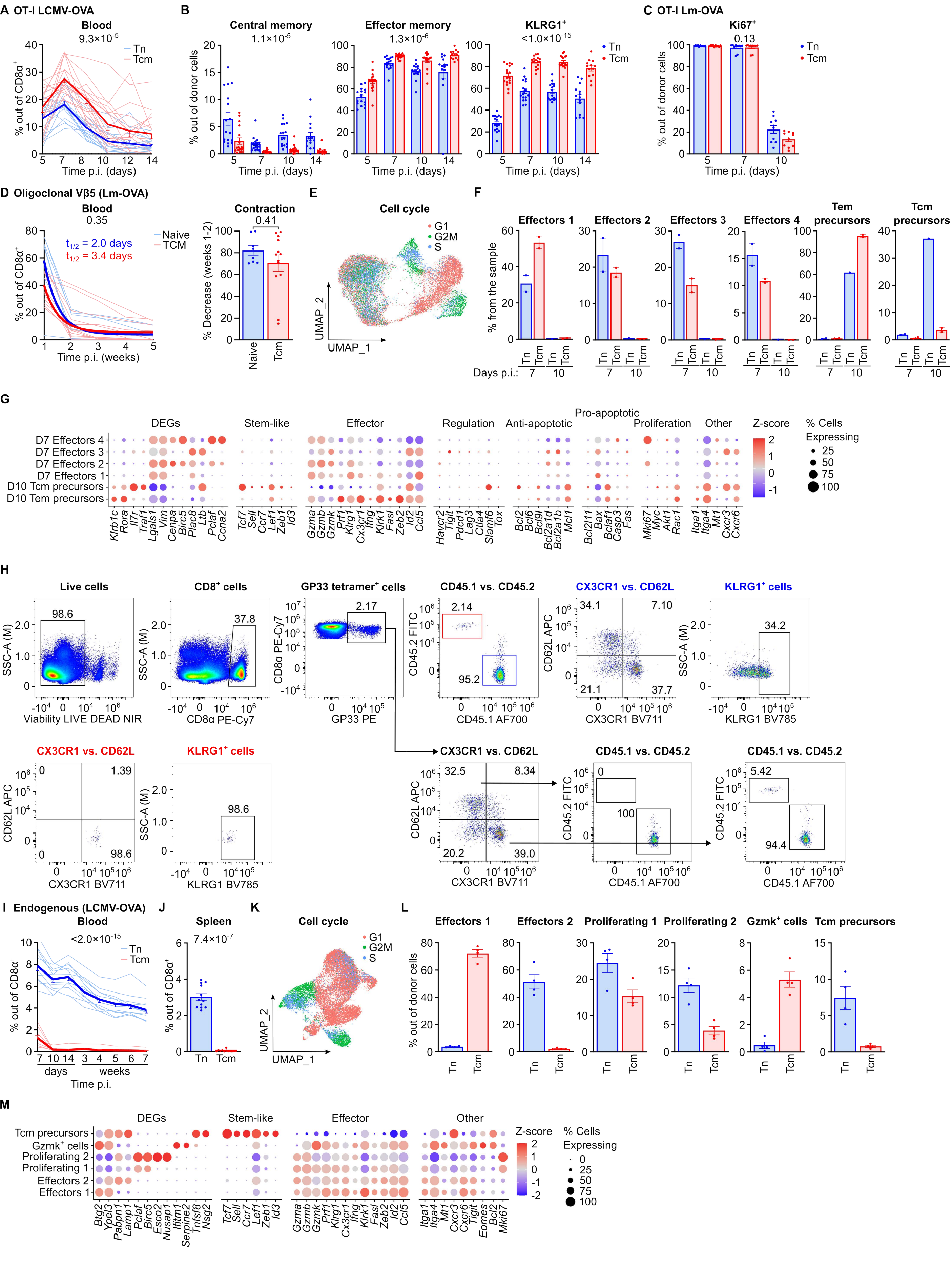
A OT-I LCMV-OVA
Blood
9.3×10-5
Tn
Tcm
% out of CD8α+
Time p.i. (days)
B
Central memory
1.1×10-5
Effector memory
1.3×10-6
KLRG1+
<1.0×10-15
% out of donor cells
Time p.i. (days)
C OT-I Lm-OVA
Ki67+
0.13
D Oligoclonal Vβ5 (Lm-OVA)
Blood
0.35
t1/2 = 2.0 days
t1/2 = 3.4 days
Naive
TCM
Time p.i. (weeks)
Contraction
0.41
% Decrease (weeks 1-2)
Naive
Tcm
E
Cell cycle
G1
G2M
S
UMAP_1
UMAP_2
F
Effectors 1
Effectors 2
Effectors 3
Effectors 4
Tem precursors
Tcm precursors
% from the sample
Days p.i.:
G
DEGs
Stem-like
Effector
Regulation
Anti-apoptotic
Pro-apoptotic
Proliferation
Other
Z-score
% Cells Expressing
D7 Effectors 4
D7 Effectors 3
D7 Effectors 2
D7 Effectors 1
D10 Tcm precursors
D10 Tem precursors
H
Live cells
CD8+ cells
GP33 tetramer+ cells
CD45.1 vs. CD45.2
CX3CR1 vs. CD62L
KLRG1+ cells
Viability LIVE DEAD NIR
CD8α PE-Cy7
GP33 PE
CD45.1 AF700
CX3CR1 BV711
KLRG1 BV785
SSC-A (M)
CD45.2 FITC
CD62L APC
I Endogenous (LCMV-OVA)
Blood
<2.0×10-15
days
weeks
Time p.i.
J
Spleen
7.4×10-7
K
Cell cycle
L
Effectors 1
Effectors 2
Proliferating 1
Proliferating 2
Gzmk+ cells
Tcm precursors
% out of donor cells
M
DEGs
Stem-like
Effector
Other
Tcm precursors
Gzmk+ cells
Proliferating 2
Proliferating 1
Effectors 2
Effectors 1

**Figure 4S**

**A-B** $2\times10^4$ Tn ($CD8^+$ $CD44^-$ ) or Tcm ($CD8^+$ $CD62L^+$ $CX3CR1^-$) were co-transferred into CD45.1/2 hosts, followed by LCMV-OVA infection. Cell progenies were monitored in the blood. n = 17 mice in 3 independent experiments. Data from one of the experiments were included in Figure 3I-J.

**A** The expansion and persistence of Tn and Tcm progeny cells in the blood were determined by their frequency among all $CD8\alpha^+$ T cells on the indicated days p.i. Thin lines show individual donor mice. The thick line represents the mean ± SEM.

**B** Phenotype of donor progenies in the blood. Samples with less than 0.1 % progenies in the blood (n = 1-4) were excluded.

**C** Percentage of $Ki67^+$ cells in the experiment shown in Figure 4C. n = 10 mice in 2 independent experiments.

**D** Expansion and persistence of oligoclonal Vβ5 $K^b$-OVA $tetramer^+$ progenies in the blood (the same experiment as in Figure 4G). Thin lines show individual donor mice. Thick lines represent fitted nonlinear regression curves (one phase decay) and the half-lives are indicated. n = 4-8 (Tn), 6-13 (Tcm) in 3 independent experiments.

**E-G** Additional analyses of the scRNA-seq experiment on progenies of oligoclonal H-$2K^b$-OVA $tetramer^+$ Tn and Tcm on day 7 or 10 p.i. (Lm-OVA) presented in Figure 4H-I.

**E** UMAP showing cell cycle phases.

**F** Frequencies of cells in particular clusters in individual mice.

**G** Dot plot showing expression of selected genes in particular clusters.

**H-M** Additional analysis of endogenous H-$2D^b$-GP33 $4\text{-mer}^+$ Tn and Tcm cells (experiment shown in Figure 4J-M).

**H** A representative gating strategy.

**I** Expansion and persistence of endogenous (Tn) and Tcm H-$2D^b$-GP33 $tetramer^+$ progeny cells in the blood following LCMV infection. Thin lines show individual donor mice. The thick line represents the mean ± SEM.

**J** Frequency of endogenous H-$2D^b$-GP33 $tetramer^+$ Tn and Tcm progenies in the spleen at week 8 p.i. (LCMV).

**K** UMAP showing cell cycle phases.

**L** Frequencies of cells in particular clusters in individual mice.

**M** Dot plot showing expression of selected genes in particular clusters.

Statistical significance was calculated using two-tailed Mann-Whitney test for a comparison of two samples (Tn vs Tcm progeny) or two-way ANOVA with mixed-effects analysis (for comparison across multiple time-points).

### Figure 5S

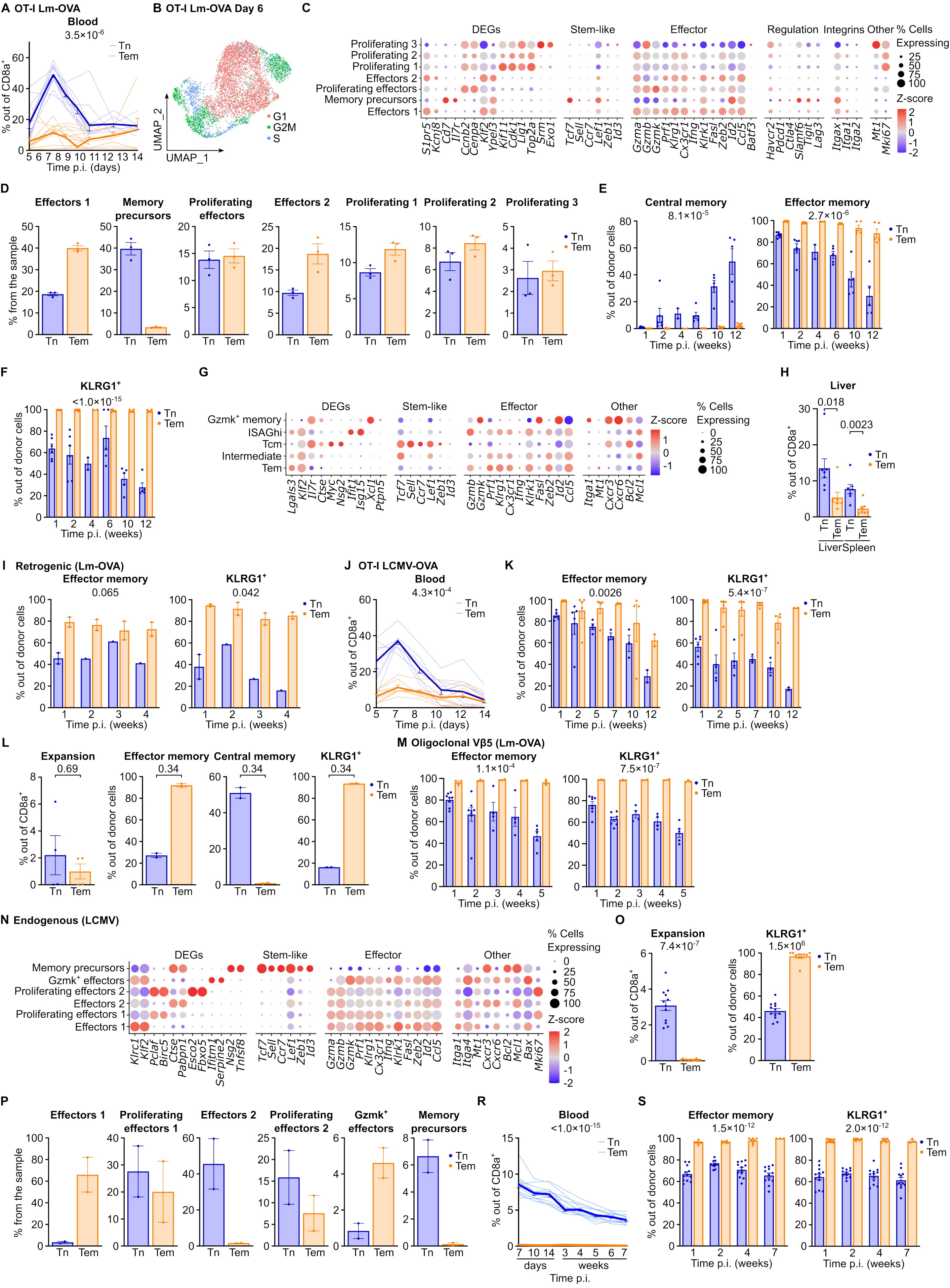
A OT-I Lm-OVA
Blood
3.5×10^-6
Tn
Tem
% out of CD8a+
Time p.i. (days)
B OT-I Lm-OVA Day 6
UMAP_1
UMAP_2
G1
G2M
S
C
DEGs
Stem-like
Effector
Regulation
Integrins
Other
% Cells Expressing
Z-score
Proliferating 3
Proliferating 2
Proliferating 1
Effectors 2
Proliferating effectors
Memory precursors
Effectors 1
S1pr5 Kcnj8 Cd7 Il7r Ccnb2 Cenpa Klf2 Ypel3 Kif11 Cdk1 Lig1 Top2a Srm Exo1
Tcf7 Sell Ccr7 Lef1 Zeb1 Id3
Gzma Gzmb Gzmk Prf1 Klrg1 Cx3cr1 Ifng Klrk1 Fasl Zeb2 Id2 Ccl5 Batf3
Havcr2 Pdcd1 Ctla4 Slamf6 Tigit Lag3
Itgax Itga1 Itga2
Mt1 Mki67
D
Effectors 1
Memory precursors
Proliferating effectors
Effectors 2
Proliferating 1
Proliferating 2
Proliferating 3
% from the sample
E
Central memory
8.1×10^-5
Effector memory
2.7×10^-6
% out of donor cells
Time p.i. (weeks)
F
KLRG1+
<1.0×10^-15
G
Gzmk+ memory
ISAGhi
Tcm
Intermediate
Tem
Lgals3 Klf2 Il7r Ctse Myc Nsg2 Ifit1 Isg15 Xcl1 Ptpn5
Tcf7 Sell Ccr7 Lef1 Zeb1 Id3
Gzmb Gzmk Prf1 Klrg1 Cx3cr1 Ifng Klrk1 Fasl Zeb2 Id2 Ccl5
Itga1 Mt1 Cxcr3 Cxcr6 Bcl2 Mcl1
H
Liver
0.018
0.0023
Liver
Spleen
I Retrogenic (Lm-OVA)
Effector memory
0.065
KLRG1+
0.042
J OT-I LCMV-OVA
Blood
4.3×10^-4
Time p.i. (days)
K
Effector memory
0.0026
KLRG1+
5.4×10^-7
L
Expansion
0.69
Effector memory
0.34
Central memory
0.34
KLRG1+
0.34
M Oligoclonal Vβ5 (Lm-OVA)
Effector memory
1.1×10^-4
KLRG1+
7.5×10^-7
N Endogenous (LCMV)
Memory precursors
Gzmk+ effectors
Proliferating effectors 2
Effectors 2
Proliferating effectors 1
Effectors 1
Klrc1 Klf2 Pclaf Birc5 Ctse Pabpn1 Esco2 Fbxo5 Ifitm1 Serpine2 Nsg2 Tnfsf8
Tcf7 Sell Ccr7 Lef1 Zeb1 Id3
Gzma Gzmb Gzmk Prf1 Klrg1 Cx3cr1 Ifng Klrk1 Fasl Zeb2 Id2 Ccl5
Itga1 Itga4 Mt1 Cxcr3 Cxcr6 Bcl2 Mcl1 Bax Mki67
O
Expansion
7.4×10^-7
KLRG1+
1.5×10^6
P
Effectors 1
Proliferating effectors 1
Effectors 2
Proliferating effectors 2
Gzmk+ effectors
Memory precursors
R
Blood
<1.0×10^-15
days
weeks
Time p.i.
S
Effector memory
1.5×10^-12
KLRG1+
2.0×10^-12

**Figure 5S**

**A** Expansion and presence of Tn and Tem progeny in blood post-Lm-OVA infection detected by FC. Thin lines show individual mice; thick lines represent means with SEMs. n = 3-12. Data from one experiment were included in Figure 5A,G.

**B** UMAP showing cell cycle stages (from Figure 5B-C).

**C** Expression of indicated genes in clusters (from Figure 5B-C).

**D** Frequencies of cells in each indicated cluster per sample (from Figure 5B-C).

**E-F** Percentage of Tem, Tcm (E), and KLRG1$^+$(F) cells among the Tn and Tem progeny over 12 weeks p.i. from the experiment in Figure 5A. Samples with less than 0.1 % progenies in the blood (n = 1) were excluded.

**G** Expression of indicated genes in clusters (additional data to Figure 5D-E).

**H** Presence of Tn and Tem cells in liver and spleen 12 weeks after Lm-OVA infection, detected by FC. Data from n = 7 across 2 experiments.

**I** Percentage of Tem and KLRG1$^+$ cells among the Tn and Tem retrogenic (Clone 2) CD8$^+$ T cell progeny over 4 weeks p.i. from the experiment in Figure 5H.

**J** Expansion and presence of Tn and Tem progeny in blood post-LCMV-OVA infection detected by FC. Thin lines show individual mice; thick lines show means with SEMs. n = 8 from 3 experiments. Data from 2 experiments shown in Figure 5I.

**K** Percentage of Tem and KLRG1$^+$ cells among Tn and Tem OT-I CD8$^+$ T cell progeny following LCMV-OVA infection over 12 weeks p.i., from the experiment in Figure 5I.

**L** The presence and percentage of Tem, Tcm, and KLRG1$^+$ cells among the Tn and Tem progeny in the spleen at 12 weeks post-LCMV-OVA infection were determined by FC at indicated time points. n = 4 mice from 2 experiments in Figure 5I. Only donor populations with at least 20 cells were included, n = 2 per group.

**M** Percentage of Tem and KLRG1$^+$ cells among Tn and Tem oligoclonal Vβ5 CD8$^+$ T cell progeny over 5 weeks p.i. from the experiment in Figure 5J.

**N** Expression of indicated genes in clusters (additional data to Figure 5L-M).

**O** Presence of Tn and Tem progeny and proportion of KLRG1-positive donor cells in the spleen at week 8 post-LCMV infection. Additional data to Figure 5N. A mouse with less than 0.1 % Tem progenies in the spleen (n = 1) was excluded from the KLRG1 proportion evaluation.

**P** Frequencies of cells within specified clusters by samples (additional data to Figure 5L-M).

**R** Expansion and persistence of Tem and endogenous H-2D$^b$-GP33 tetramer$^+$ progeny in blood following LCMV infection from Fig.5K. Thin lines show individual mice; thick lines show means with SEMs.

**S** Percentage of Tem and KLRG1$^+$ cells among Tn and Tem GP33 4-mer$^+$ CD8$^+$ T cell progeny. Additional data to Figure 5K.

### Figure 6S

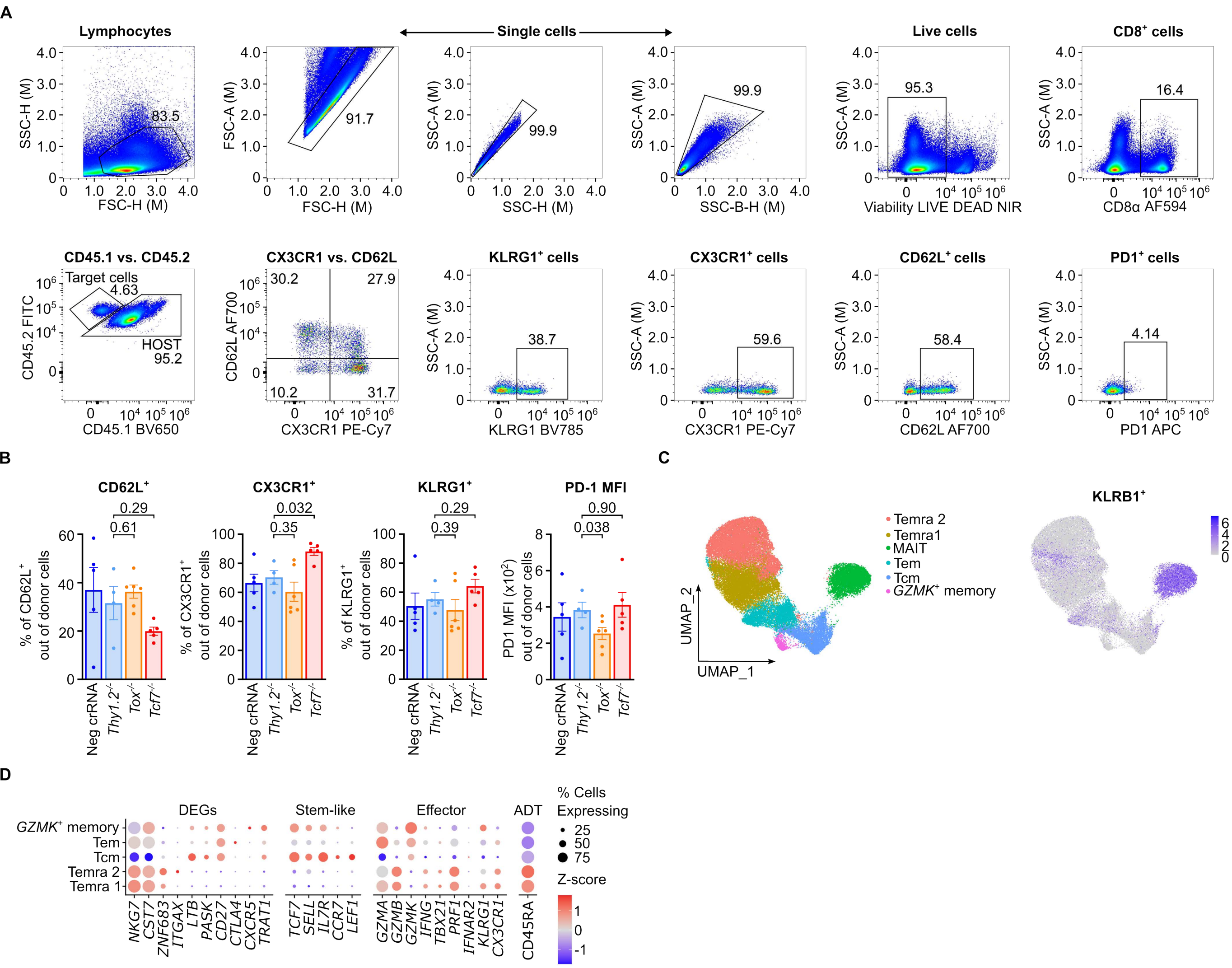
A
Lymphocytes
Single cells
Live cells
CD8+ cells
FSC-H (M)
SSC-H (M)
FSC-A (M)
SSC-A (M)
SSC-B-H (M)
Viability LIVE DEAD NIR
CD8α AF594
83.5
91.7
99.9
99.9
95.3
16.4
CD45.1 vs. CD45.2
CX3CR1 vs. CD62L
KLRG1+ cells
CX3CR1+ cells
CD62L+ cells
PD1+ cells
Target cells
4.63
HOST
95.2
CD45.2 FITC
CD45.1 BV650
CD62L AF700
CX3CR1 PE-Cy7
30.2
27.9
10.2
31.7
38.7
59.6
58.4
4.14
KLRG1 BV785
CD62L AF700
PD1 APC
B
CD62L+
CX3CR1+
KLRG1+
PD-1 MFI
0.29
0.61
0.032
0.35
0.29
0.39
0.90
0.038
% of CD62L+ out of donor cells
% of CX3CR1+ out of donor cells
% of KLRG1+ out of donor cells
PD1 MFI (x10^2) out of donor cells
Neg crRNA
Thy1.2-/-
Tox-/-
Tcf7-/-
C
Temra 2
Temra1
MAIT
Tem
Tcm
GZMK+ memory
UMAP_1
UMAP_2
KLRB1+
D
DEGs
Stem-like
Effector
ADT
% Cells Expressing
25
50
75
Z-score
GZMK+ memory
Tem
Tcm
Temra 2
Temra 1
NKG7
CST7
ZNF683
ITGAX
LTB
PASK
CD27
CTLA4
CXCR5
TRAT1
TCF7
SELL
IL7R
CCR7
LEF1
GZMA
GZMB
GZMK
IFNG
TBX21
PRF1
IFNAR2
KLRG1
CX3CR1
CD45RA

**Figure 6S**

**A** Representative gating of the data shown in Figure 6A.

**B** Frequency or mean fluorescence intensity of donor progenies positive for indicated markers in the spleen 28 p.i. (additional data to Figure 6A).

**C** UMAP plots showing unsupervised clustering before MAIT cell exclusion. MAIT cell cluster is characterized by *KLRB1* gene expression (additional data to Figure 6B-D).

**D** A dot plot of indicated genes showing differences among clusters (additional data to Figure 6C). CD45RA marker was detected by Antibody-Derived Tag (ADT).
